# Learning to suppress likely distractor locations in visual search is driven by the local distractor frequency

**DOI:** 10.1101/2022.04.29.489854

**Authors:** Fredrik Allenmark, Bei Zhang, Zhuanghua Shi, Hermann J. Müller

**Affiliations:** General and Experimental Psychology, LMU Munich, Munich, Germany; Institute of Science and Technology for Brain-Inspired Intelligence, Fudan University, Shanghai, China

**Author notes:** Correspondence concerning this article should be addressed to Fredrik Allenmark.

**Keywords:** visual search, visual attention, search guidance, attentional priority, attentional capture, statistical learning, habituation

## Abstract

Salient but task-irrelevant distractors interfere less with visual search when they appear in a display region where distractors have appeared more frequently in the past (‘distractor-location probability cueing’). This effect could reflect the (re-)distribution of a global, limited attentional ‘inhibition resource’. Accordingly, changing the frequency of distractor appearance in one display region should also affect the magnitude of interference generated by distractors in a different region. Alternatively, distractor-location learning may reflect a local response (e.g., ‘habituation’) to distractors occurring at a particular location. In this case, the local distractor frequency in one display region should not affect distractor interference in a different region. To decide between these alternatives, we conducted three experiments in which participants searched for an orientation-defined target while ignoring a more salient orientation distractor that occurred more often in one vs. another display region. Experiment 1 varied the ratio of distractors appearing in the frequent vs. rare regions (60/40–90/10), with a fixed global distractor frequency. The results revealed the cueing effect to increase with increasing probability ratio. In Experiments 2 and 3, one (‘test’) region was assigned the same local distractor frequency as in one of the conditions of Experiment 1, but a different frequency in the other region – dissociating local from global distractor frequency. Together, the three experiments showed that distractor interference in the test region was not significantly influenced by the frequency in the other region, consistent with purely local learning. We discuss the implications for theories of statistical distractor-location learning.

**Public Significance Statement:** We are frequently distracted by salient visual stimuli which are irrelevant to the task at hand. Previous studies have shown that ‘knowledge’ of the location(s) where a distractor is most likely to occur helps the observer to mitigate distraction. In this study we compared different theories of how the frequency and spatial distribution of distractor occurrence in different locations could influence the ability to avoid distraction. The results favored a local learning account: the ability to avoid distraction by distractors occuring in a particular spatial region is primarily influenced by how often distractors have occurred in that region.

## Introduction

In everyday life, we often need to ignore irrelevant information while focusing on what is important for the task at hand. However, salient events can capture our attention even when they are irrelevant. Interestingly, attentional capture by salient but irrelevant distractor stimuli can be reduced as a consequence of increased exposure to such distractors. This has been shown in ‘additional-singleton’ visual-search paradigms, in which participants search for a singleton target (defined, e.g., by a unique orientation or color) in search displays that, on some proportion of trials, also contain an additional, task-irrelevant, singleton distractor (e.g., Goschy et al., 2014; Sauter et al., 2018; Wang & Theeuwes, 2018; Zhang et al., 2019). Response times (RTs) to the target in such search tasks are generally slower on trials on which a singleton distractor is present as compared to absent – an interference effect taken as indicative of inadvertent ‘attentional capture’ by the distractor on a proportion of trials. Importantly, distractor interference can be significantly down-modulated by statistical learning. In particular, the interference is substantially reduced when distractors occur frequently as compared to only rarely anywhere in the search display (Müller et al., 2009; Won et al., 2019), where this learning effect may also exhibit a degree of position specificity: if distractors occur more frequently in one region (encompassing multiple locations) or at one specific location of the display, distractors at those locations cause less interference compared to distractors at rare locations (Allenmark et al., 2019; Goschy et al., 2014; Sauter et al., 2018; Wang & Theeuwes, 2018; Zhang et al., 2019). In analogy to the ‘target-location probability-cueing’ effect (see below), this reduction of interference for frequent distractor locations is often referred to as ‘distractor-location probability cueing’. However, while distractor-location probability cueing has been demonstrated in many studies, the underlying mechanisms remain controversial (Gaspelin & Luck, 2018a; Geng et al., 2019; van Moorselaar et al., 2021; van Moorselaar & Slagter, 2020).

Preceding the recent focus on distractor handling, RTs and response accuracy in search tasks had been established to depend on the probability distribution of target locations: RTs are faster and response accuracy is improved when the target on a given trial appears at a location where it had been encountered more frequently in the past (e.g., Geng & Behrmann, 2002; Shaw & Shaw, 1977). In classical demonstration of this target-location probability-cueing effect (e.g., Geng & Behrmann, 2002; Shaw & Shaw, 1977), participants searched for a single, briefly presented, letter (shown alone, without any non-targets, in the display for brief, 20–30-ms durations and terminated by a noise mask), which appeared more often at some locations than others, and reported whether the letter was a ‘T’, ‘E’, or ‘V’. Shaw and Shaw (1977) showed that the accuracy in this target task could be well explained by a model which assumed that a limited, attention-related resource was optimally allocated (in terms of maximizing the overall target-detection probability) based on the frequency distribution of the location of the target letter. By analogy, distractor-location probability cueing might similarly reflect optimal distribution of a limited resource, such as assigning less of a (target-)attention-related resource or more of a distractor-inhibition-related resource to frequent versus rare distractor locations.

Critically, according to an optimal resource-distribution account, distractor-location probability learning would be global, in the sense that changing the spatial distribution of distractor occurrence would influence not only the amount of interference at any given location, but also the interference at any other locations – because allocating a greater portion of a limited resource to one location reduces the portion available for the other locations. Alternatively, distractor-location probability learning might be a purely local effect, such that the frequency with which a distractor occurs at one location influences only that specific location (and perhaps some nearby locations). One possible local mechanism could be ‘habituation’, which has been proposed to play a role in learning to filter out irrelevant distractors (Duncan & Theeuwes, 2020; Turatto, Bonetti, & Pascucci, 2018; Turatto, Bonetti, Pascucci, et al., 2018; Won & Geng, 2020). In the context of distraction paradigms, habituation would mean that the attentional ‘orienting reflex’ to a distractor stimulus becomes weaker with increased exposure to the stimulus (Sokolov, 1963). However, while habituation could be local, it could conceivably also be global, or a combination of both (e.g., Valsecchi & Turatto, 2021).

Habituation could be local if repeated exposure to distractors in a spatial region results in a reduced ‘orienting reflex’ to, and consequently reduced interference by, distractors in that region. It could also be global if observers learn to ignore objects with certain features regardless of where they occur, that is: repeated exposure to a particular type of distractor (e.g., a display item singled out from the non-distractor items by a particular color, such as a *red* shape among *green* shapes) may lead to reduced interference from such (red) distractors regardless of where they occur in the display (consistent with reports that in search for a shape-defined target, the interference caused by singleton color distractors decreases with globally increasing distractor frequency; e.g., Bogaerts et al., 2022; Geyer et al., 2008; Müller et al., 2009; Won et al., 2019). Such global habituation would necessarily be (distractor-) feature- (or, more generally, dimension-) based, whereas local habituation may be purely location-based, or local-feature based, or a combination of both. Of note, only local habituation could explain (locally) reduced distractor interference for a frequent, compared to a rare, distractor region or location. Also, global habituation makes the opposite prediction to the limited-resource account in terms of global effects, namely: globally reduced distractor interference even if distractor frequency is increased in a specific region or location, versus reduced interference in that region or location along with increased interference by distractors occurring at other locations. The aim of the present study was to examine these differential predictions regarding global effects in order to distinguish between the different local and global theories of distractor-location probability cueing.

In order to distinguish between the different local and global accounts of distractor-location probability cueing outlined above, we performed three visual search experiments. In all experiments, the task was to find an orientation-defined target, a bar tilted 13° from the vertical, while ignoring a distractor defined in the same dimension as the target (orientation), a horizontally oriented bar, where this distractor appeared more often in the top or the bottom region of the search display (see Figure 2 below). Compared to the more widely used ‘different-dimension distractors’ (such as a color singleton in search for a shape-defined target), distractors defined in the same dimension as the target give rise to greatly increased interference effects not only in terms of RTs, but also oculomotor and electrophysiological measures of true attentional capture (i.e., mis-guidance of the eye or covert attention to the distractor location, prior to re-allocation of the eye or attention to the target; e.g., Sauter et al., 2018, 2021; Liesefeld, Liesefeld, & Müller, 2017). Along with the greater capture effects, same-dimension distractors also generate increased distractor-location probability-cueing effects compared to different-dimension distractors (Sauter et al., 2018, 2021; Liesefeld & Müller, 2021). We systematically varied the frequency with which (same-dimension) distractors occurred in the frequent and rare distractor regions, along with (in some cases) the global distractor frequency, between experiments or between groups within an experiment.

**Figure 1:**
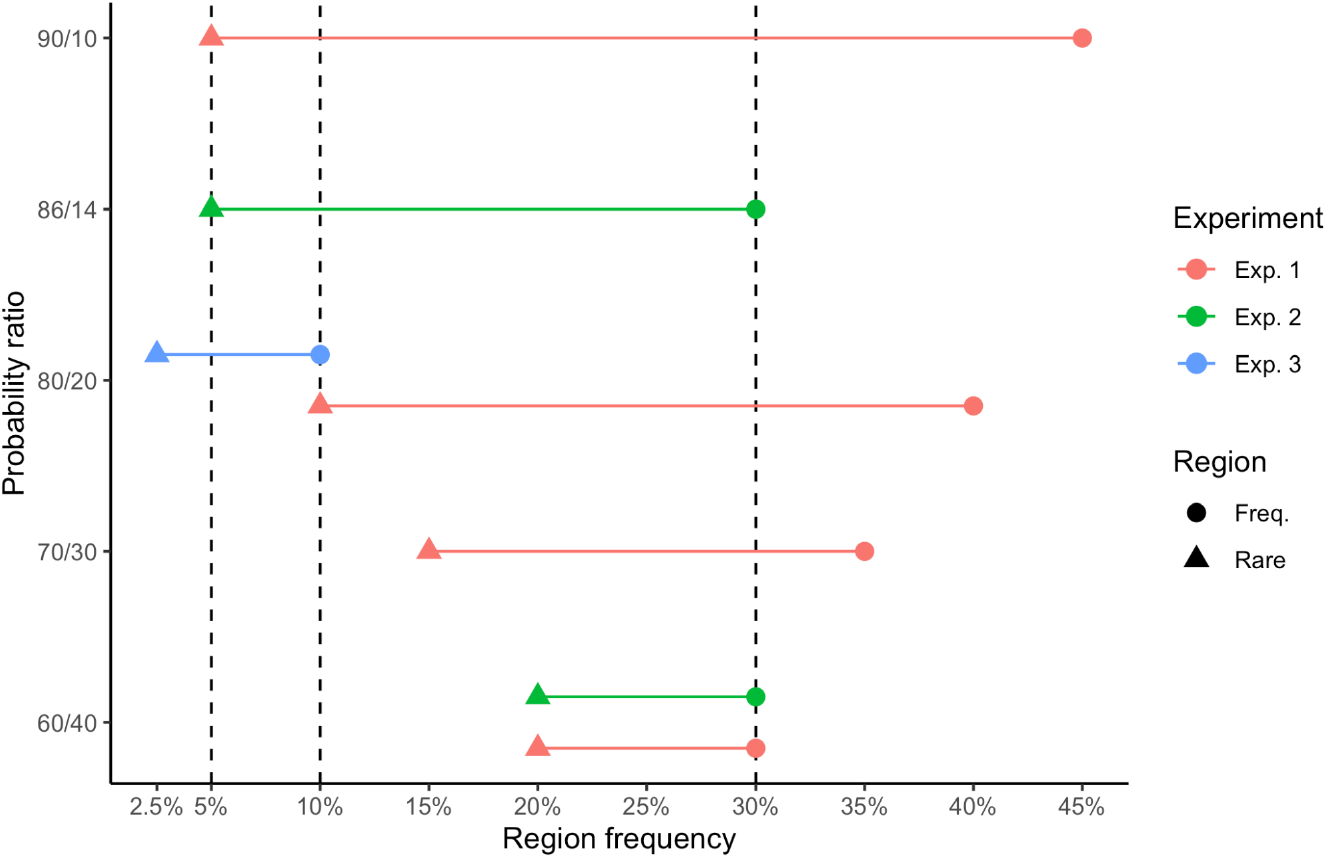
Visual illustration of the different experimental conditions performed by the different groups in the three experiments (listed in Table 1). Each pair of dots connected by a line indicates the frequent and rare distractor region in one experimental group. The x-coordinates represent the region distractor frequency, while the y-coordinates represent the ratio between the frequent and rare region distractor frequencies in a group. Critically, the vertical dashed lines indicate the same distractor region frequency across experiments. The local habituation account would predict the same distractor interference for the same region frequency.

**Figure 2:**
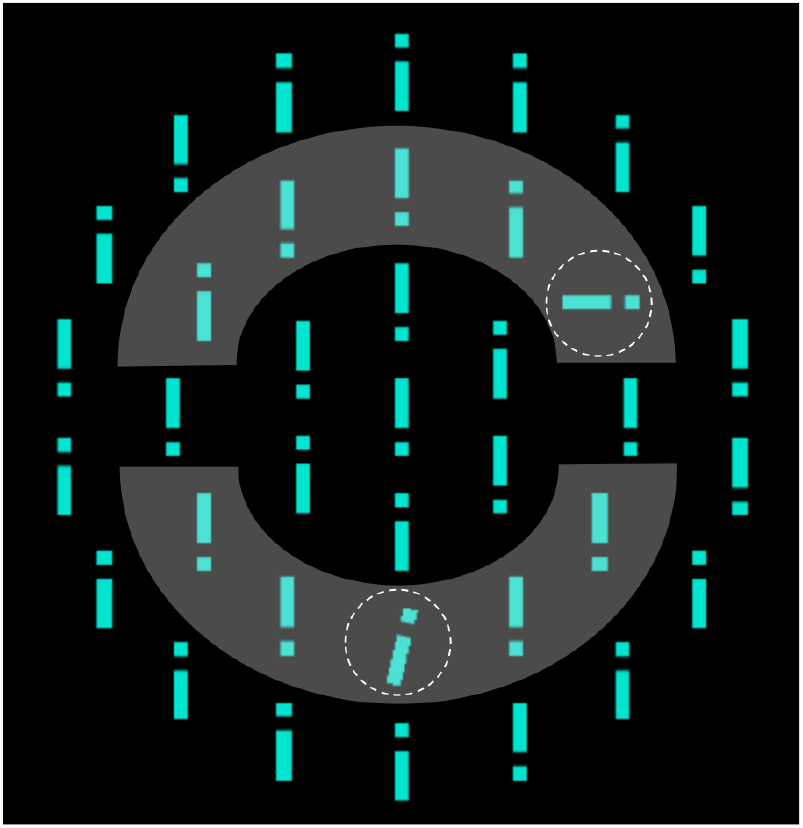
Example of a search display, with a singleton distractor (top right, outlined by the dashed circle) and a target (at the bottom of the middle ring, outlined by the dashed circle). Participants responded based on the location of the dot on the i-shaped target, which in this case is on the top. The target and the distractor appeared only in the middle ring gray regions. The upper and lower gray-shaded semicircle regions indicate the frequent and, respectively, rare distractor regions. Note the dashed circles and gray area are for illustration purposes only; they were not shown in the experiment.

The aim of Experiment 1 was to examine how the reduction of distractor interference in a frequent compared to a rare region depends on the distractor probability ratio between the two regions, while keeping the global distractor frequency (i.e., singleton distractor occurrence) fixed. To this end, we tested, in different groups of participants, four different conditions, all with a distractor in half of the trials, but with different ratios of distractors in the frequent and rare regions: 60/40, 70/30, 80/20, and 90/10, respectively. Both the local-habituation and the limited-resource theory predict that distractor interference in the frequent distractor region should be the lower and the interference in the rare region the higher the more unequal the ratio. Thus, Experiment 1 was mainly intended to provide a baseline against which to compare the results of the other two experiments.

Experiment 2 was designed to examine the global effects, if any, of changing the distractor frequency in one region – specifically by reducing the distractor frequency in the rare region by 75% compared to the 60/40 condition of Experiment 1 –, while keeping the distractor frequency in the other – the frequent – region fixed (thereby also reducing the overall distractor frequency; see Figure 1). In this case, the different theories make different predictions of how, if at all, distractor interference in the *frequent* region (where the distractor frequency was unchanged) would differ from the 60/40 condition Experiment 1: (i) ‘local habituation’ predicts no difference in distractor interference since only the local frequency matters; (ii) the ‘limited-resource’ account predicts reduced interference since the frequent region should receive more of the limited resource by lowering the frequency of distractors in the rare region; and (iii) ‘global habituation’ would predict increased interference since the global distractor frequency is reduced.

**Table 1:** The different experimental conditions performed by the different groups in the three experiments. The distractor region ratio column shows the ratio of the distractor presence in the frequent vs. rare regions. The column of “Distractor prevalence” indicates the percentage of distractor-present trials in the total trials, while the columns of distractor region frequency showed the regional frequencies (frequent vs. rare) of the distractor-present trials.

| Group | Distractor ratio | Distractor Prevalence | Distractor region frequency |  |
| --- | --- | --- | --- | --- |
|  |  |  | Frequent | Rare |
| E1 60/40 | 60 / 40 | 50% | 30% | 20% |
| E1 70/30 | 70 / 30 | 50% | 35% | 15% |
| E1 80/20 | 80 / 20 | 50% | 40% | 10% |
| E1 90/10 | 90 / 10 | 50% | 45% | 5% |
| E2 50% | 60 / 40 | 50% | 30% | 20% |
| E2 35% | 86 / 14 | 35% | 30% | 5% |
| E3 | 80 / 20 | 12.5% | 10% | 2.5% |

Finally, Experiment 3 had two purposes, the first being to examine the effect of global distractor frequency, while keeping the ratio between the two regions fixed: we used the ratio of 80/20, which had been tested in one condition in Experiment 1, but with global distractor prevalence reduced from 50% (in Experiment 1) to 12.5% (in Experiment 3). Additionally, Experiment 3 allowed us to examine whether the amount of interference caused by distractors in a region depends on whether this region is the frequent or the rare distractor region, since the frequent region in Experiment 3 was designed to have the same local distractor frequency as the rare region in the 80/20 condition of Experiment 1. (i) The limited-resource theory would predict that, even though the local frequency is the same, distractor interference should be higher in the rare region in the 80/20 condition of Experiment 1 compared to the frequent region in Experiment 3; (ii) ‘global habituation’ would make the opposite prediction (because distractors are less frequent overall in Experiment 3 compared to Experiment 1); and (iii) ‘local habituation’ would predict no difference in distractor interference between the rare region in the 80/20 condition of Experiment 1 and the frequent region in Experiment 3 (because the local frequency is the same).

The whole study was designed as a one coherent whole (to realize all necessary comparisons to decide among the three accounts) to compare conditions within a given experiment but also across experiments. However, due to the restrictions on laboratory studies imposed by the CORONA pandemic, only one experiment could be conducted onsite (Experiment 1), while two had to be run online (Experiments 2 and 3), as participants earmarked for these experiments could no longer be invited to the laboratory. To ensure comparability across experiments, we ran conditions with the same distractor-region ratio (60/40) and a distractor prevalence of 50% both onsite (Experiment 1) and online (Experiment 2). The two experiments yielded similar results, in terms of both the common, distractor-absent (trial) baseline and the distractor-interference magnitude on trials with a distractor in the frequent and, respectively, the rare region.

More generally, the planned cross-group comparisons may be compromised by different groups by chance exhibiting different levels of baseline performance (RTs on distractor-absent trials). For instance, one group may process the search displays generally slower than another, magnifying their distractor-interference effects. Accordingly, a difference in interference may be falsely attributed to the manipulation of the local/global distractor probability, when in fact it just reflects a baseline shift. To guard against this, and following previous studies (e.g., Brascamp et al., 2011; Kruijne et al., 2015), we also examined the patterns of interference effects related (‘normalized’) to the distractor-absent baseline. Of note, the essential results turned out just the same whether examined in terms of the absolute or the normalized interference scores.

To preview the outcome of the study: overall, the data argue strongly in favor of ‘local’ learning of, and adaptation to, the frequencies with which distractors occur at particular display locations (relatively independent of the global distractor frequency). In the Discussion, we consider the relation of local learning to ‘habituation’ accounts of the interference reduction engendered by statistical distractor-location learning, as well as where, in the functional architecture of search guidance, local learning may be implemented.

## Experiment 1

Experiment 1 examined how the amount of interference caused by distractors occurring in a frequent and, respectively, a rare distractor region changes as a function of the bias in the distractor-location distribution, importantly with a fixed global distractor frequency.^1^ For this purpose we compared four bias conditions, tested in four different groups of participants. In each condition, a distractor was present in half of the trials and was more likely to occur in one versus the other region of the search display (either top or bottom region, counterbalanced across participants), with the proportion of distractors that appeared in the frequent region varying from 60% to 90% (see Figure 1).

### Methods

#### Participants

A total of 88 participants (mean age: 26.7 years; 60 females) were recruited from the student population at Ludwig-Maximilians-University (LMU) Munich, with 22 participants randomly assigned to each of the four groups. The sample size was determined based on previous studies, in particular, Sauter et al. (2018), who – with same-dimension distractors and a probability ratio of 90/10 between the frequent and rare distractor region – had an effect size of dz=1.3 for the distractor-location probability-cueing effect. Proceeding from this, we chose the sample size to have 90% power with half this effect size (assuming a one-tailed test since we expected distractor interference to be reduced, not increased, in the frequent distractor region), in order have enough power to resolve the smaller distractor location effects that we expected to find in the conditions with smaller probability ratios. In addition, we wanted to ensure sufficient power for resolving the differences in the size of the probability-cueing effect among (at least the more extreme of) the four different probability-ratio groups. A previous study by Lin et al. <u>(2021)</u>, which used a very similar between-group design to ours, achieved an effect size of 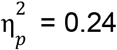 for the critical interaction between the within-participant factor Distractor Location (frequent, rare location) and the between-participant factor (frequent/rare-location) Probability Ratio. Assuming this effect size, we calculated that we would have more than 99% power for our comparison of the probability-cueing effect across groups (the actual effect size turned out to be somewhat larger: 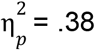; see results section below).

The study protocol was approved by the Ethics Committee of the LMU Faculty of Psychology and Pedagogics. Informed consent was obtained from all participants prior to the experiment. They were remunerated at a rate of 9 Euro per hour for their service.

#### Apparatus

The experiment was conducted in a moderately lit test room. Stimuli were presented on a 24″ LCD monitor with a 1920 × 1080 pixels screen resolution. Stimuli were generated by Psychophysics Toolbox Version 3 (PTB-3) (Brainard, 1997) based on MATLAB (The MathWorks Inc.). Participants viewed the monitor from a distance of 60 cm (eye to screen) and gave their responses by pressing the keys “y’ and ‘m’, with their left and right index fingers respectively, on a QWERTZ keyboard.

#### Stimuli

The stimuli were essentially the same as those we had used in several previous studies (e.g., Sauter et al., 2018, 2019, 2020; Zhang et al., 2021). The search displays consisted of 37 gray bars, which, except for one central bar, were arranged around three concentric circles (2°, 4°, and 6° of visual angle in radius), presented on a black background (see Figure 2). We used dense displays, with multiple rings, in order to ensure that local feature contrast was sufficiently high to result in pop-out of the singleton target and distractor items (see Liesefeld et al., 2016). Each bar was 0.25° in width and 1.35° in length. Further, each bar contained a gap, 0.25° in size, located 0.25° from either the top or the bottom of the bar at random (forming an ‘i’- or inverted ‘i’-type stimulus). All bars were vertical except the tilted singleton target (13° clockwise or counter-clockwise from the vertical), which was present on all trials, and a horizontally oriented singleton distractor, present in 50% of trials. Targets and distractors occurred only on the middle (4°) ring and never in the left- and right-most positions on that ring (because these lie on the horizontal midline and so do not belong to either the ‘top’ or ‘bottom’ parts of the display). Participants’ task was to find the target and respond to the position of the gap in this bar (top vs. bottom, i.e., “i” vs. inverted “i”).

#### Design

The four groups of participants performed the same search task, the only difference being the frequency distribution of the distractor locations (Figure 1). All groups performed 1440 trials, divided into 12 blocks of 120 trials. The target appeared equally frequently in both (the top and bottom) halves of the search display. By contrast, the distractor appeared more often in one (either the top or bottom) half of the search display (the frequent distractor region), compared to the other half (the rare distractor region). We used a design with a frequent distractor region, rather than a single frequent location, in order to avoid confounds between distractor- and target-location probability (see Zhang et al. (2019) for a discussion of why using a single frequent distractor location results in such confounds). The proportion of distractors which occurred in the frequent region differed between the groups, ranging from 60% to 90% (see Figure 1). Which half of the search display was the frequent and which was the rare region was counterbalanced across participants within each group.

#### Procedure

Each trial started with a central fixation cross, presented for a random duration between 700 and 1100 ms. This was followed by the search display, which remained on the screen until the participant responded. Participants were instructed to search for the 13°-tilted target bar (“i”) and respond, as quickly and accurately as possible, based on whether the gap in the “i” was on the top or the bottom. Responses were made by pressing the “y” or, respectively, the “m” key, on a “QWERTZ” keyboard, with stimulus-response mapping counterbalanced across participants. An error message (“Incorrect”) was displayed for 500 ms when a participant made an incorrect response. Between trial blocks, participants had the opportunity to take a break of a self-determined length. Of note, participants were not informed about the global, 50% distractor frequency and the probability ratio with which distractors appeared in the frequent and the rare region. At the end of the experiment, participants completed a questionnaire, designed to determine whether they had become aware of the frequent distractor region. They were first asked whether they thought the distractor had appeared equally often in all parts of the search display or more frequently in one region. Regardless of the response to the first question, they were then given a forced choice question with the four alternatives that distractors occurred more often in the top, bottom, left or right area of the search display.

#### Bayes-factor analysis

Bayesian analyses of variance (ANOVAs) and associated post hoc tests were carried out using JASP 0.15 (http://www.jasp-stats.org) with default settings. All Bayes factors for ANOVA main effects and interactions are inclusion Bayes factors calculated across matched models. Inclusion Bayes factors compare models with a particular predictor to models that exclude that predictor. That is, they indicate the amount of change from prior inclusion odds (i.e., the ratio between the total prior probability for models including a predictor and the prior probability for models that do not include it) to posterior inclusion odds. Using inclusion Bayes factors calculated across matched models means that models that contain higher-order interactions involving the predictor of interest were excluded from the set of models on which the total prior and posterior odds were based. Inclusion Bayes factors provide a measure of the extent to which the data support inclusion of a factor in the model.Bayesian *t* tests were performed using the ttestBF function of the R package BayesFactor with the default setting (i.e., rscale = medium).

### Results

For all RT analyses, we excluded trials on which a participant made an incorrect response. In addition, trials with RTs slower than 3 seconds or faster than 150 ms were considered as outliers and also excluded (approximately 1% of trials). Finally, the first block (120 trials) was excluded because in this block, the participant would not yet have fully learned the distractor distribution.

#### Baseline RTs

As the distractor-absent trials were equally frequent (and exactly the same) in all groups, this condition served as a common baseline against which distractor-interference and distractor-location probability-cueing effects in and among the various groups can be assessed. The average RTs (and the associated standard errors) on distractor-absent trials were 769 ± 33, 748 ± 29, 840 ± 33, and 759 ± 34 ms for the four distractor-location probability-ratio (60/40, 70/30, 80/20, 90/10) groups, respectively. A one-way ANOVA with Group as between-subject factor revealed the distractor-absent RTs to be comparable across the four groups (non-significant Group effect, *F*(3, 84) = 1.79, *p* = 0.16, 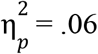, *BF*_incl_ = 0.42). Error rates were also comparably low, at around 2–3% for all groups.

#### Distractor interference and Probability-cueing effect

Given the absence of significant differences in the baseline RTs among the groups, we went on to examine the distractor-interference effects on response speed, that is, the difference in RTs between trials with and without a distractor. Figure 3A depicts the interference caused by distractors in the frequent and, respectively, the rare region as a function of the probability ratio. As can be seen, interference in the frequent region tended to decrease with increasing probability ratio (though this trend was not statistically significant: *F*(3, 84) = 2.15, *p* = .10, BF_incl_ = 0.61), whereas interference in the rare region tended to increase (*F*(3, 84) = 3.41, *p* < .05, 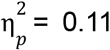, *BF*_incl_ = 2.3); Bonferroni-corrected post-hoc *t*-tests indicate the interference to be significantly larger in the 80/20 compared to the 60/40 group (*t*(42) = 3.11, *p*_bonf_ < .05, *BF*_10_ = 12.6).

**Figure 3:**
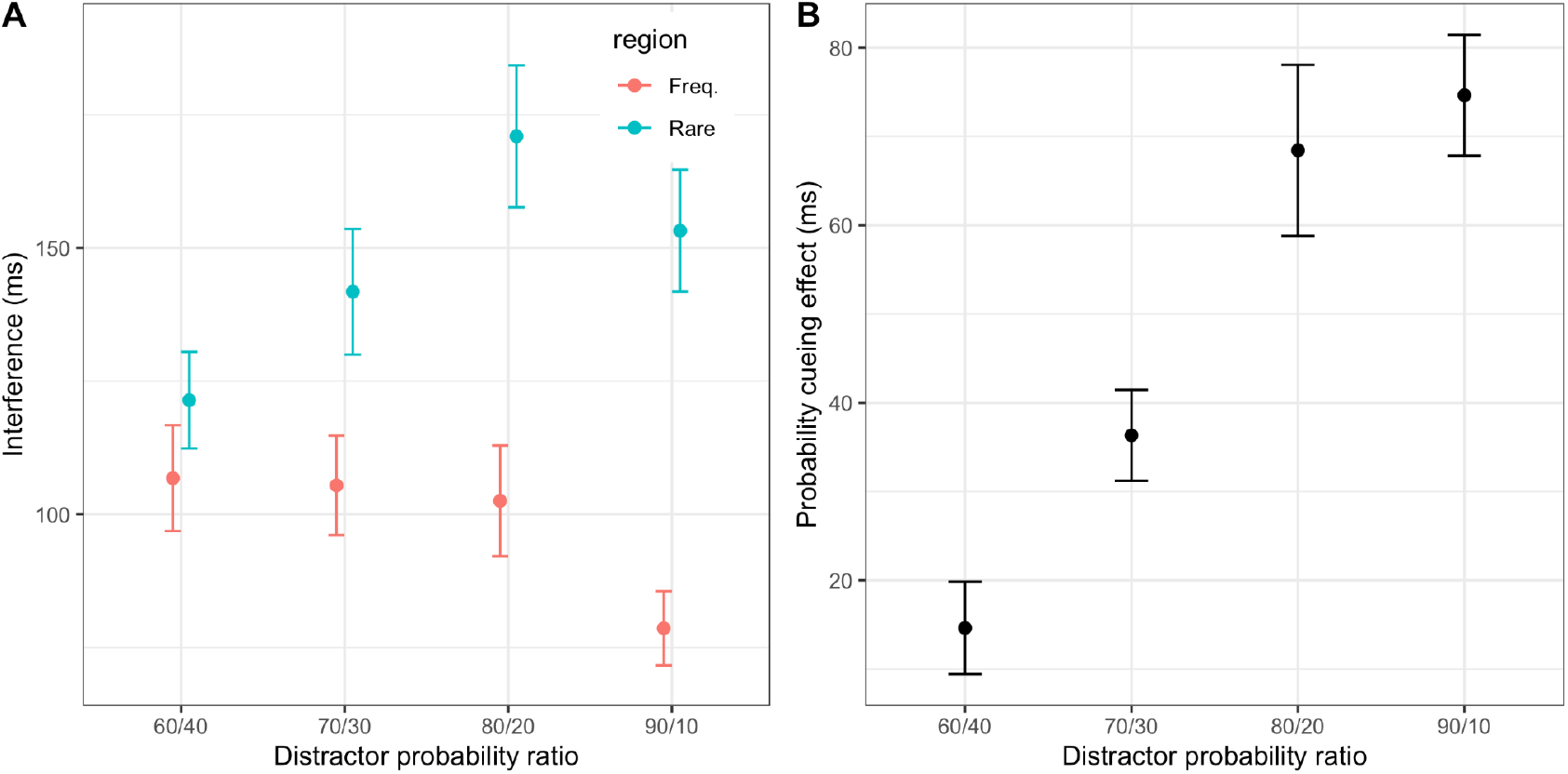
On the top left is the distractor-interference effect on RT (A, top row), calculated as the RT difference between trials with a distractor compared to distractor-absent trials, when a distractor occured in the frequent region (red) or in the rare region (cyan). On the top right is the distractor-location probability-cueing effect on RT (B, top row), defined as the RT difference between trials with a distractor in the rare vs. the frequent distractor region. The bottom row shows the corresponding effects on the error rates. Error bars indicate the standard error of the mean.

The difference in interference between the two regions (Figure 3A) is the distractor-location probability-cueing effect, which is depicted in Figure 3B as a function of of the different distractor-distribution groups. As can be seen, the probability-cueing effect increased with increasing probability ratio of the distractor distribution (*F*(3, 84) = 17.2, *p* < .001, 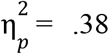, *BF*_incl_ > 1000). Bonferroni-corrected post-hoc tests showed that the probability-cueing effect was significantly larger in the 90/10 condition compared to both the 70/30 condition and the 60/40 condition, and significantly larger in the 80/20 condition compared to both the 70/30 condition and the 60/40 condition (all *t*s(42) > 3.3, *p*s_bonf_ < .01, BFs_10_ > 9.4). Numerically the probability cueing effect was also larger in the 90/10 compared to 80/20 condition and in the 70/30 compared to 60/40 condition, but these differences were not significant after correcting for multiple comparisons (*p*s_bonf_ > .15).

#### Target-location effect

Figure 4 presents the target-location effect, that is, the difference in mean RTs between trials on which the target appeared in the frequent versus the rare distractor region. As can be seen, the target-location effect was generally positive (i.e., RTs were slower to targets in the frequent vs. the rare region) and increased with increasing probability ratio between the two distractor regions, for each distractor condition (distractor absent, distractor in rare region, and distractor in frequent region). A mixed-effects ANOVA yielded a significant main effect of Group (i.e., Probability Ratio) (*F*(3, 84) = 4.2, *p* < .01, 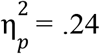, *BF*_incl_ = 5.7); Bonferroni-corrected post-hoc tests revealed the target-location effect to be significantly larger in the most biased, 90/10, versus the least biased, 60/40, condition (*t*(42) = 3.3, *p*_bonf_ = .009, *BF*_10_ > 1000). Of note, the interaction between Group and Distractor Condition was non-significant (*F*(6, 168) = 0.69, *p* = .65, *BF*_incl_ = 0.04), and the Bayes factor strongly favors models without an interaction term – indicating that the effect of the probability ratio on target processing was largely independent of whether or not, and in which region a distractor appeared in the display. There was, however, a significant main effect of distractor condition (*F*(2, 168) = 26.0, *p* < .001, 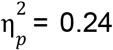, BF_incl_ > 1000): the target-location effect was overall largest on trials with a distractor in the frequent region (59 ms compared to 15 ms with a distractor in the rare region and 25 ms on distractor-absent trials). This could reflect an effect of proximity between the distractor and the target: with a distractor in the frequent region, targets in the frequent region would be closer to the distractor, potentially slowing RTs on these trials compared to trials with a target in the rare region.

**Figure 4:**
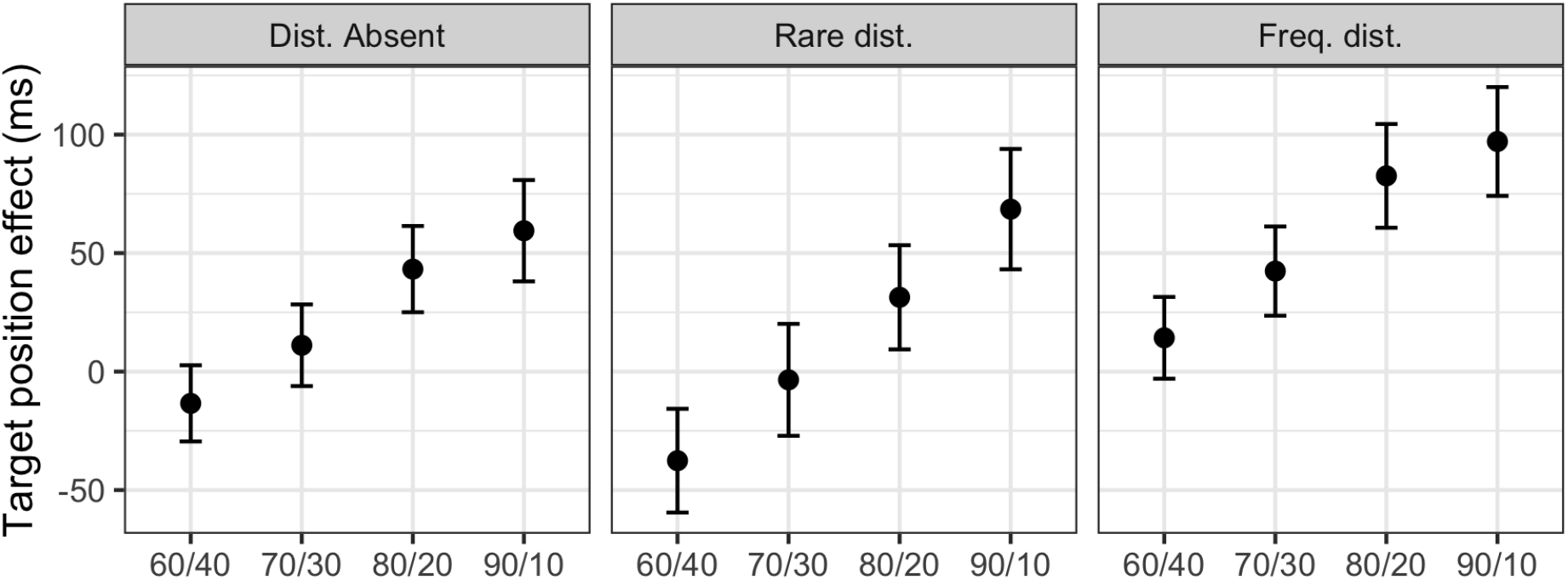
Target-location effect, i.e., difference in RT between trials with a target in the frequent vs. the rare distractor region. The left, middle, and right panels show the target-location effect on trials without a distractor, with a distractor in the rare region (Rare dist.), and with a distractor in the frequent region (Freq. dist.), respectively. Error bars indicate the standard error of the mean.

#### Awareness effects

In order to examine to what extent participants had become aware of the spatial distractor distribution, we analyzed how many of them had noticed a spatial bias in the distractor distribution and how many indicated the frequent distractor region correctly in the subsequent forced-choice recognition test (see Methods for details). In the groups with the 60/40, 70/30, 80/20, and 90/10 probability ratios, 41%, 55%, 46%, and, respectively, 50% of participants believed the distractor distribution had been biased. And the proportion of participants who answered the forced-choice question correctly was 41%, 36%, 46%, and 50% – which was larger than expected by chance (i.e., 25%). Binomial tests revealed this difference to be significant in the two groups with the largest probability ratios (*p* = .044 for the 80/20 group and *p* = .012 for the 90/10 group). But note that of those who believed the distribution was biased, only 44%, 33%, 60%, and 46% indicated the frequent region correctly.

To check whether the distractor-probability cueing effect differed between participants who had vs. had not correctly indicated the regularities, we performed an ANOVA on the cueing effect with Probability Ratio and Awareness (correct vs. incorrect recognition response) as factors. While the main effect of Awareness was non-significant and the Bayes factor favors models without an awareness effect (*F*(3, 80) = 0.04, *p* = .85, *BF*_incl_ = 0.23), Awareness interacted significantly with the Probability Ratio (*F*(3, 80) = 3.63, *p* < .05, 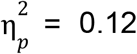, *BF*_incl_ = 3.5): this interaction was due to aware participants exhibiting a larger cueing effect in the 60/40 and 80/20 probability-ratio conditions but a smaller effect in the 70/30 and 90/10 conditions. However, comparing the data collapsed across the two lower and, respectively, the two higher probability ratios, there was no difference between the ‘aware’ (cueing effects of 24.5 vs. 71.5 ms) and the ‘unaware’ (cueing effects of 26.0 vs. 73.0 ms) group – suggesting that the interaction reflects a chance finding.

### Discussion

Overall, the results of Experiment 1 revealed a trade-off in distractor interference between the frequent and rare regions: the larger the probability ratio was (i.e., the more/less distractors appear in the frequent/rare region), the less the interference caused by a distractor in the frequent region and the more the interference by a distractor in the rare region, that is: the larger the statistical-learning or ‘distractor-location probability-cueing’ effect. This effect was mirrored in a target-location effect: RTs were generally slower to targets in the frequent (vs. the rare) distractor region, with this difference growing with the probability ratio of distractors appearing in the frequent/rare region. This pattern would be consistent with some limited inhibitory resource being divided between (or distributed across) the frequent and rare distractor regions according to the probability ratio. However, it would also be consistent with ‘habituation’ accounts, which would predict habituation to distractors to be the greater (and, thus, distractor interference to be the smaller) the more frequently distractors occur in a given region. Experiments 2 and 3 were designed to differentiate between these two types of account.

## Experiment 2

Experiment 2 was designed to compare the global ‘limited-resource’ account and ‘local habituation’ account. The global ‘limited-resource’ would predict that the amount of distractor interference in the frequent region would depend on the frequency in the rare region, importantly, even when the frequency in the frequent region is fixed, while the ‘local habituation’ account predicts that the frequency of distractors in the rare region should make no difference to interference in the frequent region. For this reason, we manipulated the distractor prevalence (one group with 50% and the other 35%) such that the distractor region frequency was the same in the frequent region, but differed in the rare region (see Table 1).

Due to restrictions related to the COVID-19 pandemic, we performed this experiment as an online experiment. We programmed the experiment in PsychoPy and ran it on the online platform Pavlovia (www.pavlovia.org). In order to ensure that any differences compared to the 60/40 condition of Experiment 1 were not due to changes introduced to run the experiment online, we also replicated the 60/40 condition of Experiment 1 as an online experiment.

### Methods

#### Participants

A total of 2 x 22 participants (Group 1, replication of 60/40 condition in Experiment 1: mean age: 25.3 years; 9 females; Group 2, reduced-frequency group: mean age: 26.2 years; 12 females) were recruited for this experiment. Each participant was paid 15 Euro in compensation. Informed consent was obtained from all participants prior to the experiment.

#### Apparatus and Stimuli

To equate the conditions as well as possible with the onsite (laboratory) setup in Experiment 1, participants were told to perform the (online) experiment in a quiet, and moderately lit, environment, on full screen (laptop or external monitor). The display monitor was to be placed on a table surface, with the participant being seated on a chair, with their hands comfortably resting on the (response) keyboard in front of them and viewing the monitor at arm’s length (i.e., a distance of some 60 cm); and the display brightness was to be set to a middle contrast.

Since Experiment 2 was conducted as an online experiment, each participant ran it on their own computer, with potentially quite different monitor sizes. In order to ensure comparable conditions across the different participants, participants were asked to enter their monitor size into the participant information box at the beginning of the experiment. All stimuli were then scaled according to the respective monitor size. In addition, we instructed them to view the monitor from an arm’s length distance (approximately 60 cm), so as to also keep the stimulus size in degrees of visual angle comparable across participants (and to Experiment 1). – In all other respects, the search displays were the same as those used in Experiment 1.

#### Design and Procedure

Two groups performed different conditions, differing in terms of the proportion of trials on which a singleton distractor was present and the frequency distribution of the distractor position (see Figure 1). In each group, the distractor appeared more often in one half (either the top or bottom half, counterbalanced across participants) of the search display (the frequent distractor region), compared to the other half (the rare distractor region). One group was a replication of the condition with the least biased (60/40) distribution of the distractor across the frequent and rare regions in Experiment 1 (in which the distractor prevalence was 50%, with 30% and 20% of distractors appearing in the frequent and rare regions, respectively). In the other group, the distractor prevalence was reduced to 35% and the distractor ratio increased to 86/14 to keep the frequency with which distractors appeared in the frequent region the same (30% of all trials had a distractor in the frequent region) as the other group. However, the frequency with which distractors occurred in the rare region was reduced by a factor of four (from 20% to 5% of trials) – In all other respects, the task and trial procedure was the same as in Experiment 1.

### Results and Discussion

As for Experiment 1, outlier trials with RTs slower than 3 seconds or faster than 150 ms (approximately 1% of trials), as well as trials on which an incorrect response was made, were excluded from RT analysis. In addition, the first block (120 trials), during which participants would gradually learn the distractor-probability distribution, was excluded.

Figure 5 presents the main results of Experiment 2, compared to the 60/40 condition in Experiment 1. Since one condition in (onsite) Experiment 2 was a replication of the 60/40 condition in (onsite) Experiment 1 (both with a distractor prevalence of 50% and distractor-location probability ratio of 60/40), we first compared this condition to the corresponding condition of Experiment 1, which yielded no significant differences between the online and onsite experiments in the average distractor-absent RTs: *t*(42) = 0.78, *p* = .44, *BF*_10_ = 0.38 or in the distractor-location probability-cueing effect: *t*(42) = 0.79, *p* = .43, *BF*_10_ = 0.38. We further performed a mixed-effect ANOVA for the target-location effect with the factors Experiment (online and onsite) and Distractor Condition (distractor absent, distractor in frequent region, distractor in rare region), which again revealed no significant effects involving Experiment (main effect, *F*(1, 42) = 1.22, *p* = .27; Experiment x Distractor-Condition interaction (*F*(2, 84) = 0.68, *p* = .51). However, the main effect of Distractor Condition was significant (*F*(2, 84) = 14.5, *p* < .001): the target-location effect was largest on trials with a distractor in the frequent region (as already observed in Experiment 1).

**Figure 5:**
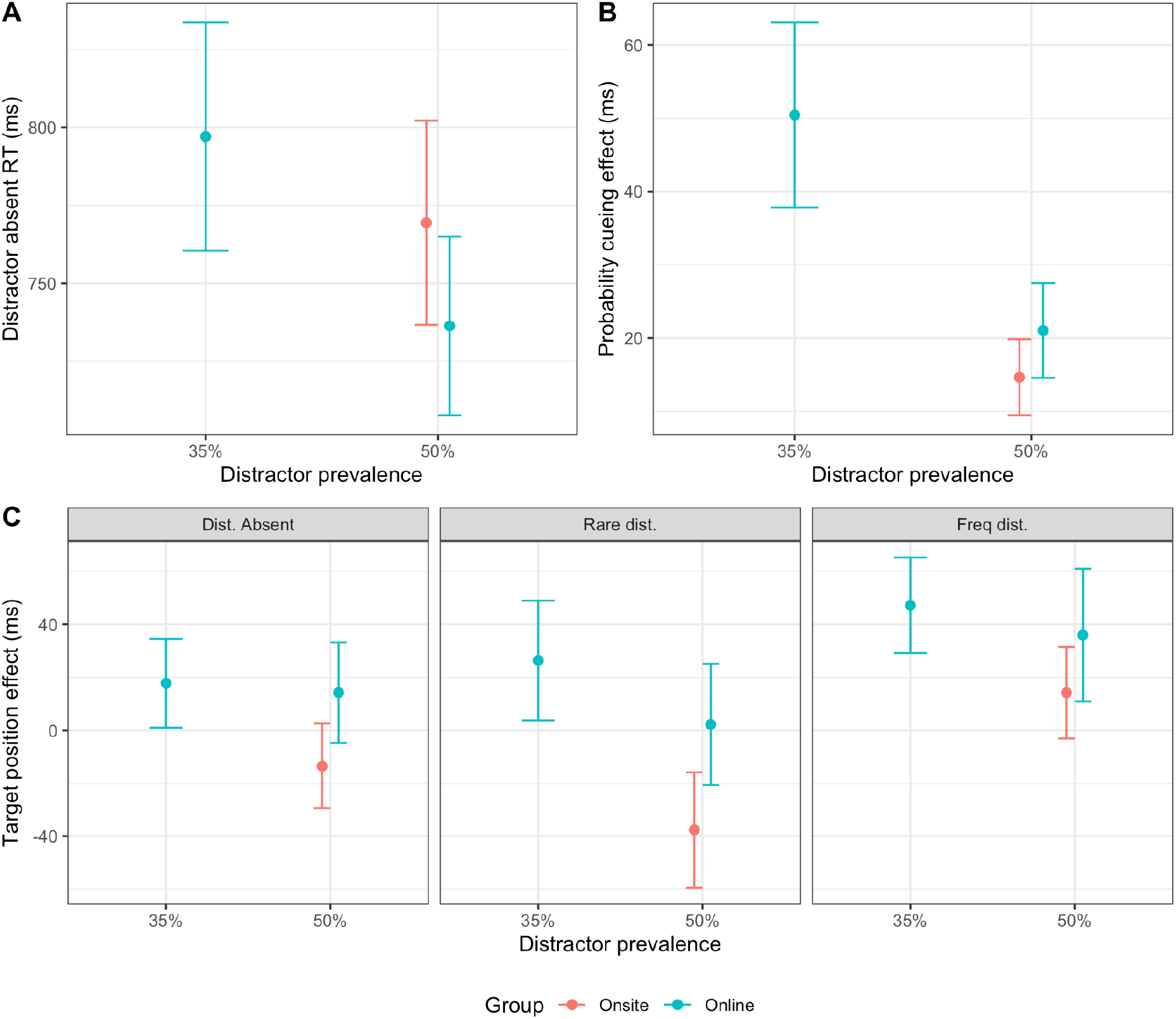
Summary of results for online Experiment 2 (35% and, respectively, 50% distractor prevalence), and the corresponding 60/40 condition of onsite Experiment 1 (50% global distractor frequency): (**A**) distractor-absent RTs, (**B**) distractor-location probability-cueing effect, (**C**) and target-location effect. Error bars indicate the standard error of the mean.

Having confirmed that the online 60/40 condition replicated the main results of the 60/40 condition of onsite Experiment 1, we next compared this condition to the novel condition in Experiment 2 – which we refer to as the ‘reduced distractor-frequency condition’, as the distractor frequency in the rare region was reduced by a factor of 4 compared to the 60/40 condition, resulting in a distractor prevalence of 35% and a distribution with 86% of distractors appearing in the frequent region (see Figure 1). The average distractor-absent RTs did not differ significantly between the 60/40 condition and the reduced distractor-frequency condition (*t*(42) = 1.3, *p* = .19, *BF*_10_ = 0.61), ensuring comparable baselines.

Importantly, concerning the question at issue: the distractor-location probability-cueing effect turned out to be larger, by about 30 ms, in the reduced distractor-frequency vs. the 60/40 condition (*t*(42) = 2.1, *p* < .05, *BF*_10_ = 1.7). This is consistent with the finding in Experiment 1 that the probability-cueing effect increases with more biased distractor distributions. Given the increased distractor-location probability-cueing effect, one would have expected the target-location effect to be also larger in the reduced distractor-frequency vs. the 60/40 group. However, although the target-location effect was numerically somewhat greater in the reduced distractor-frequency group (31 ms vs 18 ms), the modulation by Group was not significant: Group main effect, *F*(1,42) = 0.23, *p* = .63; Group x Distractor-Condition interaction (*F*(2,84) = 0.61, *p* = .55). However, as before, the target-location effect turned out again largest on trials with a distractor in the frequent region: Distractor-Condition main effect, *F*(2,84) = 5.2, *p* < .01.

Additionally, we examined the distractor interference separately for trials on which distractors appeared in the frequent and the rare region (see Figure 6). Because the rare region in the reduced distractor-frequency condition of Experiment 2 had the same local (region) distractor frequency (i.e., the frequency out of all, distractor-present and -absent, trials with which a distractor appeared in that region) as the rare region in the 90/10 condition in Experiment 1, we also included the latter condition in the comparison.

**Figure 6:**
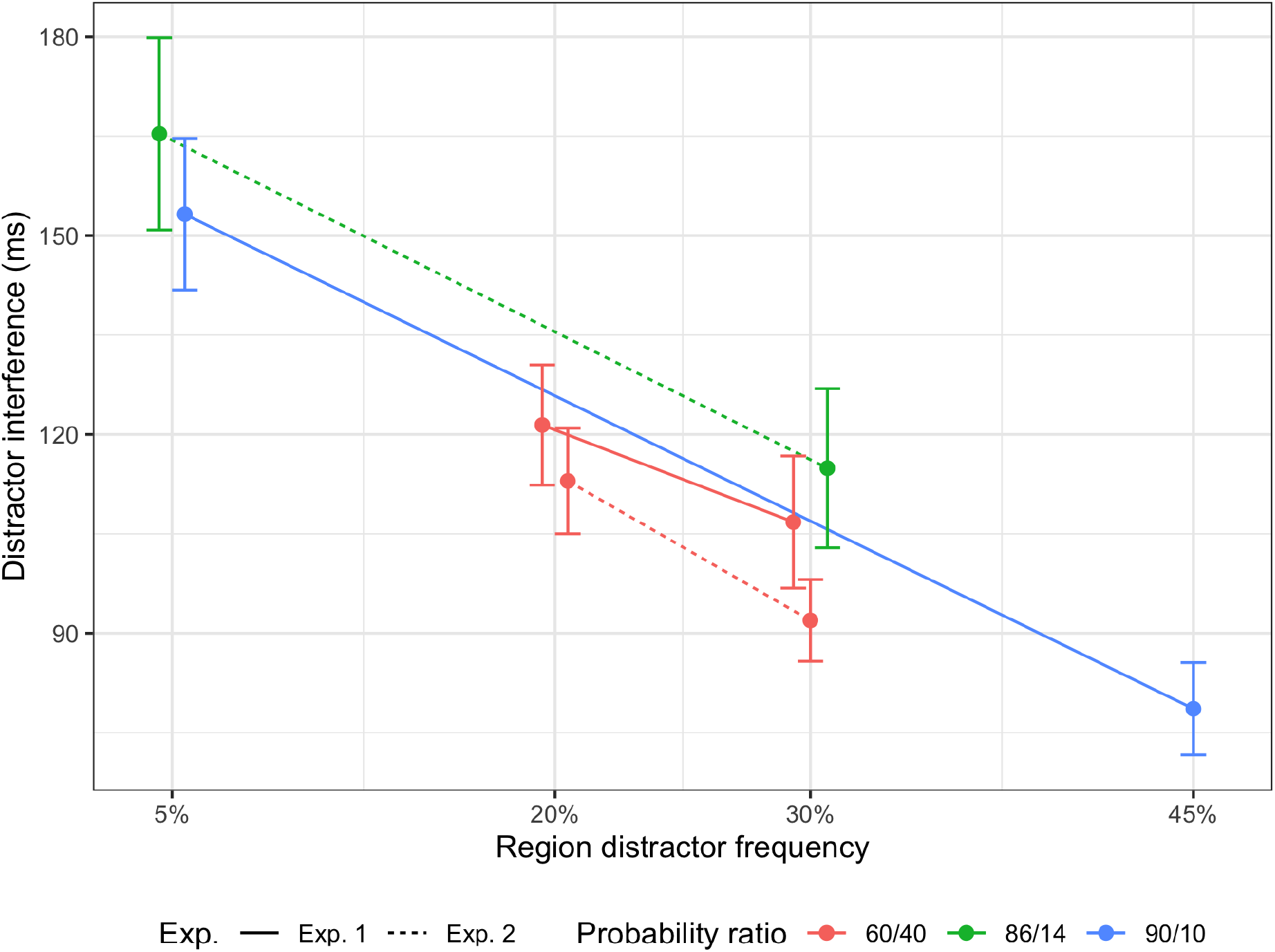
Average distractor interference as a function of the local (region) distractor frequency. Plotted are overlapping frequent and/or rare region conditions Experiment 2 (60/40 and 86/14 probability-ratio groups) and Experiment 1 (60/40 and 90/10 probability-ratio groups). Error bars indicate the standard error of the mean.

The interference caused by distractors in the rare region was significantly greater, by 52 ms, in the reduced distractor-frequency condition compared to the 60/40 reference condition of Experiment 2 (*t*(42) = 3.24, *p* < .01, *BF*_10_ = 16), as well as the equivalent, 60/40 condition of Experiment 1 (44 ms difference; *t*(42) = 2.63, *p* < .05, *BF*_10_ = 4.3). In the rare region, the local frequency of distractors was lower in the reduced distractor-frequency condition, so the increased interference is consistent with both global and local accounts of distractor-location probability cueing. Importantly, for the frequent region, distractor interference in the reduced distractor-frequency condition was neither significantly different compared to the 60/40 reference condition of Experiment 2 (23 ms difference; *t*(42) = 1.74, *p* = .09, *BF*_10_ = 0.99), nor compared to the equivalent, 60/40 condition of Experiment 1 (8 ms difference; *t*(42) = 0.53, *p* = .60, *BF*_10_ = 0.33). Since the local distractor frequency in the frequent region was the same in the different conditions, this result is consistent with local theories of distractor probability cueing, but not with global theories since the frequency in the rare region differed. Moreover, the interference caused by distractors in the rare region did not differ significantly between the reduced distractor-frequency condition of Experiment 2 and the 90/10 condition of Experiment 1 (12 ms difference, *t*(42) = 0.67, *p* = 0.50, *BF*_10_ = 0.36). Again, this is consistent with local theories of distractor probability cueing, because the rare region had the same local frequency in both conditions; but it is inconsistent with global theories, since the frequency in the other region was different. In Appendix 3, we confirm that the above three comparisons are also non-significant, with two of the Bayes factors favoring the null hypothesis and one inconclusive, when comparing normalized distractor interference (normalized by the mean distractor-absent RT for each participant). Distractor interference in the frequent region, on the other hand, was significantly larger in the reduced distractor-frequency condition of Experiment 2 compared to the 90/10 condition in Experiment 1 (36 ms difference, *t*(42) = 2.68, *p* < .05, *BF*_10_ = 4.7), as expected based on both local and global accounts.

Finally, we analyzed the responses to the post-experiment questions, testing whether participants had become aware of the biased distractor-location distribution (see Methods section for Experiment 1). In the online replication of the 60/40 condition of Experiment 1, 18% (4 out of 22) of participants responded that they thought distractors had occurred more often in one region. When forced to indicate which display half had contained a distractor more frequently, 32% (7 out of 22) of participants selected the correct region – which, while numerically somewhat larger than expected based on guessing (25%), did not differ significantly from chance level (binomial test: p = .46). Of the 7 participants who thought distractors had occurred more often in one region, only 2 chose the correct region in the forced-choice question. While the probability-cueing effect was numerically smaller for participants who responded correctly to the forced-choice question (5 ms vs. 29 ms), the difference was non-significant (Welch’s t-test: *t*(10.7) = 1.79, *p* = .10, *BF*_10_ = 1.29). they thought distractors had occurred more often in one region and 50% answered the forced-choice question correctly, the latter being significantly above chance level (binomial test: p = .012). This indicates that at least in the reduced frequency group, for which the probability ratio between the two distractor regions was larger (86/14 compared to 60/40), some participants may have become aware of the bias in the distractor distribution. However, there were only four participants who both responded that they thought distractors had occurred more often in one region and chose the correct region in the forced-choice question. Further, while the probability-cueing effect was numerically larger for participants who responded correctly to the forced-choice question (60 ms vs. 41 ms), the difference was not significant (Welch’s t-test: *t*(19.7) = 0.79, *p* = .44, *BF*_10_ = 0.48).

Thus, there was little evidence in either group that ‘awareness’ of the spatial distractor bias systematically influenced the cueing effect.

## Experiment 3

The purpose of Experiment 3 was to investigate the effect of global distractor prevalence while keeping the probability ratio between the frequent and rare distractor region fixed, as well as whether the amount of interference caused by distractors in a region depends on whether the region is the frequent or the rare distractor region. For this purpose we tested a condition in which the distractor prevalence was reduced to 12.5% as compared to 50% in Experiment 1, but the distractor-location probability-ratio remained 80/20 as in the 80/20 condition in Experiment 1. With such settings we were able to reduce the local distractor probability in the frequent region in Experiment 3 to 10%, having the same local distractor probability as in the rare region of the 80/20 condition in Experiment 1 (see Figure 1). This would allow us to explicitly test the ‘global’ vs. ‘local’ habituation accounts.

Due to restrictions related to the covid-19 pandemic, we again performed this experiment as an online experiment. Since we found no significant difference between the results of the online replication of the 60/40 condition and the original onsite experiment, we compared the results of the online condition in Experiment 3 directly to the “80/20” condition of Experiment 1, rather than replicating the later as an online experiment.

### Methods

#### Participants

A total of 22 participants (mean age: 26.5 years; 12 females) were recruited for this experiment. Each participant was paid 15 Euro as compensation. Informed consent was obtained from all participants prior to the experiment.

#### Design, Stimuli, and Procedure

Experiment 3 introduced the same, 80/20 probability ratio as in the 80/20 condition of Experiment 1, but the frequency of distractors was reduced by a factor of four in each region – yielding a global distractor frequency of 12.5, as compared to the 50% frequency in Experiment 1 (see Figure 1). The experiment consisted of a single session with 1440 trials, divided into 12 blocks of 120 trials. – In all other respects, the search displays, task, and trial procedure were the same as in Experiments 1 and 2.

### Results

All reaction time (RT) analyses excluded outlier trials with RTs slower than 3 seconds or faster than 150 ms (approximately 2% of trials) as well as trials on which a participant made an incorrect response. In addition, the first block (120 trials) was excluded because in this block the participant may not yet have learned the distractor distribution.

Because Experiment 3 was designed to have the same proportion of distractors in the frequent versus the rare region, as well as the same frequency of distractors in one of the regions, as the 80/20 group of Experiment 1, but with a different global distractor prevalence (12.5% versus 50% in Experiment 1), the results from Experiment 3 are compared to this group in the Figures 7 and 8 and analyses below.

**Figure 7:**
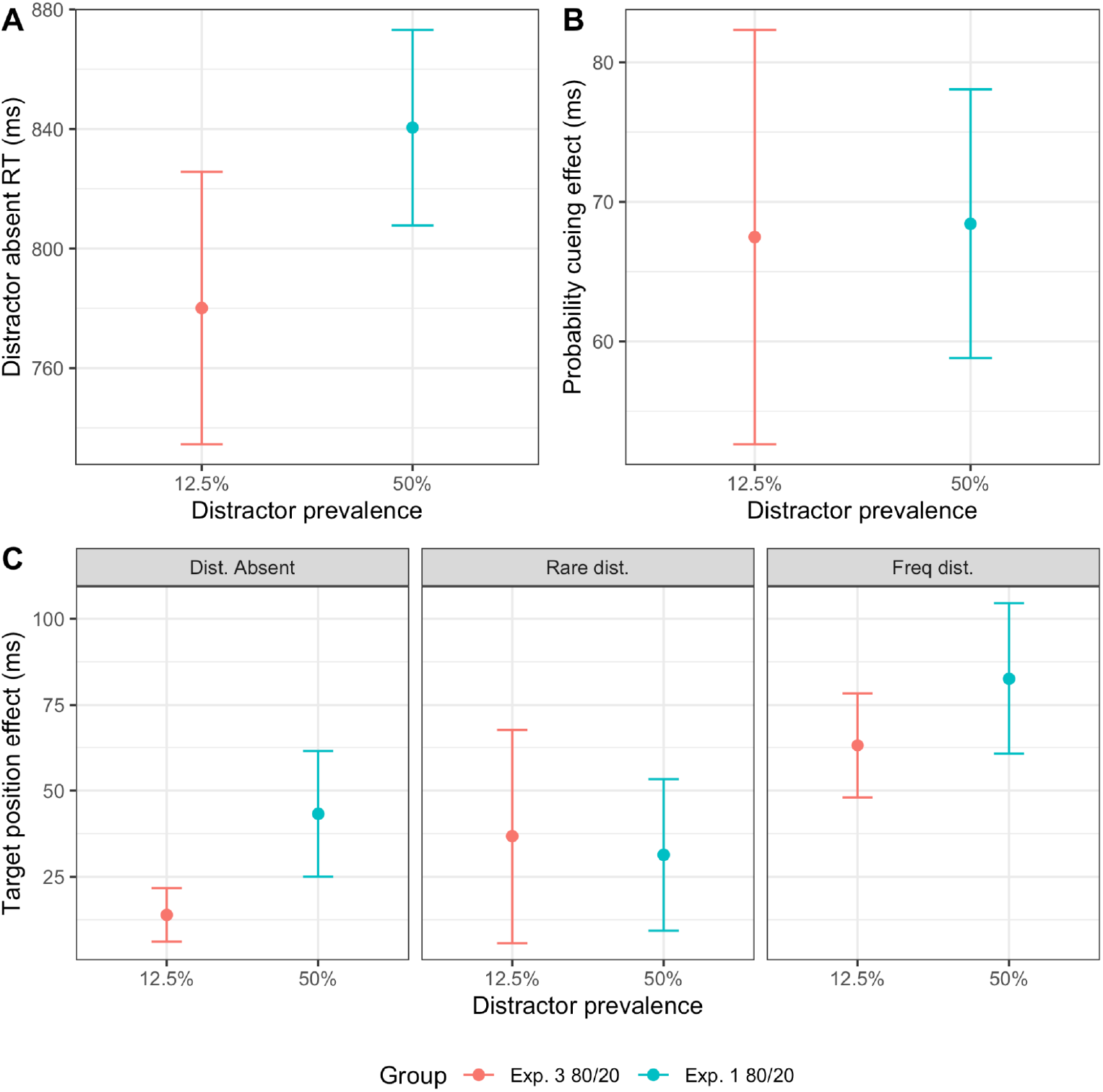
Summary of results for Experiment 3 (Distractor prevalence of 12.5%) and the corresponding 80/20 condition of Experiment 1 (Distractor prevalence of 50%): (**A**) mean RTs of the distractor-absent trials, (**B**) The probability-cueing effect and (**C**) the target-location effects, separated for the conditions of the distractor absent, distractor at rare, and respectively at the frequent regions. Error bars indicate the standard error of the mean.

**Figure 8:**
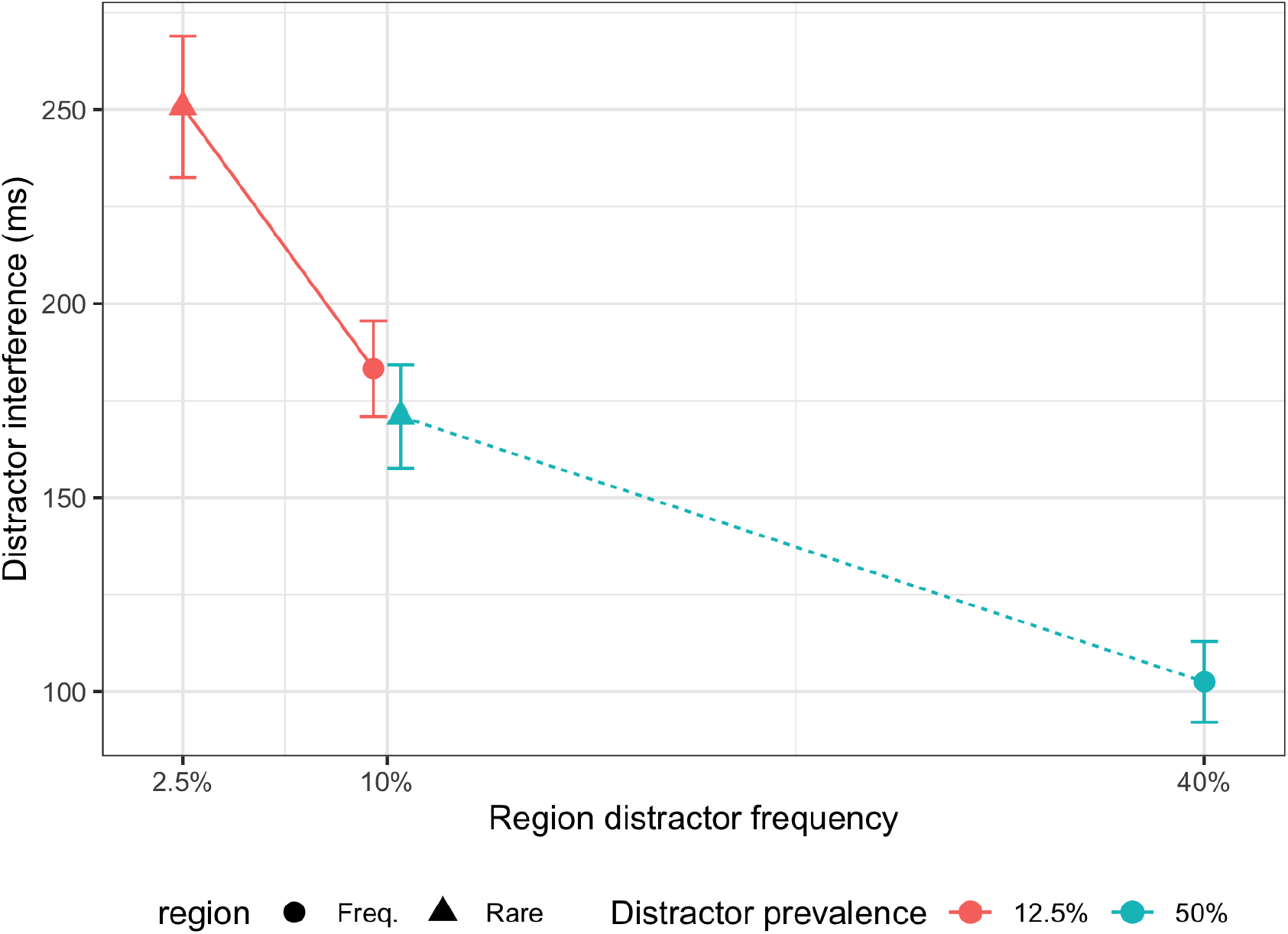
Average distractor interference on those trials where a distractor appeared in the frequent or the rare region in Experiment 3 and the corresponding “80/20” condition in Experiment 1 as a function of the frequency of distractors in the region as well as the global distractor frequency in the experimental group. Error bars indicate the standard error of the mean.

Figure 7 shows some of the main results of Experiment 3 compared to the 80/20 group of Experiment 1, which both had the same proportions of distractors in the frequent and rare regions (80% in frequent region), but differed in terms of the global distractor prevalence (50% in Experiment 1, 12.5% in Experiment 3). The mean RT of the distractor-absent trials was numerically, but not significantly faster with the lower global distractor prevalence (780 vs. 840 ms; *t*(42) = 1.1, *p* = .28, *BF*_10_=0.48). Most importantly, the distractor-location probability-cueing effect did not differ significantly between the low and high distractor-prevalence groups, with the Bayes factor supporting the null hypothesis (*t*(42) = 0.05, *p* = .96, *BF*_10_ = 0.30). While the target-location effect was numerically somewhat smaller in the group with the lower global distractor prevalence (38 ms compared to 52 ms), the difference was not statistically significant (Group main effect, *F*(1,42) = 0.38, *p* = .54; Group x Distractor-Condition interaction (*F*(2,84) = 0.80, *p* = .45). However, again it was the largest on trials with a distractor in the frequent region (Main effect of distractor condition, *F*(2,84) = 5.9, *p* < .01).

Finally, we examined the distractor interference separately for trials on which distractors appeared in the frequent and the rare region (see Figure 8). Distractor interference was higher in the group with the lower global distractor prevalence, both for distractors in the frequent region (*t*(42) = 5.1, *p* < .001, *BF*_10_ > 1000) and distractors in the rare region (t(42) = 3.6, p < .001, BF_10_ = 38). Because the frequency with which distractors occurred in the frequent region in Experiment 3 was (designed to be) the same as the frequency with which distractors occurred in the rare region in the 80/20 condition of Experiment 1, we also compared distractor interference between these two conditions. This comparison revealed no difference in distractor interference between the frequent (Experiment 3) and the rare (Experiment 1) region with the same local distractor frequency (*t*(42) = 0.69, *p* = .49, *BF*_10_ = 0.36). In Appendix 3, we confirm that this comparison is also non-significant, with a Bayes factor favoring the null hypothesis, based on normalized distractor interference. This is consistent with ‘local’, but inconsistent with ‘global’, theories of distractor-location probability cueing.

In addition, we analyzed the responses to the awareness questionnaire for the reduced distractor prevalence group. 55% (12 out of 22) of participants responded that they believed distractors had occurred more often in one region, and 55% responded correctly to the forced-choice question, which is significantly more than expected by chance (binomial test: p = .005). However, of those who answered the forced-choice question correctly, only 8/12 had indicated that they believed distractors had occurred more often in one region. And the distractor-location probability-cueing effect did not differ significantly between participants who responded correctly vs. incorrectly to the forced-choice question (Welch’s t-test: t(17.5) = 0.68, p = .50, BF_10_ = 0.46). Thus, again, there was no evidence that the cueing effect was influenced by ‘awareness’ of the spatial distractor bias.

## Within-region analysis

Across three experiments, we consistently found that interference from distractors within a defined sub-region of the search display was smaller in those groups and conditions in which the local distractor frequency in that region was higher. This was the case both within each experiment (witness the distractor-location probability-cueing effects) as well as between experiments (see, e.g., the comparison between the 80/20 probability-ratio conditions in Experiment 1 and Experiment 3 in Figure 8). Critically, however, distractor interference in a region was not significantly influenced by the distractor frequency outside of that region (see Figure 6 and Figure 8). In fact, it was not even significantly influenced by whether locations outside the region had a higher or lower distractor frequency compared to locations inside the region – as can be seen from comparing distractor interference in the rare region in the 80/20 condition of Experiment 1 and the frequent region in Experiment 3 (see Figure 8), both of which had the same local distractor frequency (a distractor appeared in the region in 10% of trials), but in one case the other region had a four times higher frequency (Experiment 1) while in the other case it had a four times lower frequency (Experiment 3). Given this, it appears justified to consider all conditions in all three experiments together in order to look for a functional dependence of distractor interference on the local distractor frequency within a given region.

However, while the interference effect did not differ significantly between those different conditions (in different experiments) that had the same local distractor frequency, numerically there was a consistent pattern of interference being (numerically) larger in the condition with the lower global distractor frequency (see Figures 6 and 8), and the Bayes factors were in some cases inconclusive. The limited-resource theory would have predicted a difference in the opposite direction, since lower global distractor frequency means lower frequency of distractors outside the region – thus, more of the limited resource should be allocated to locations inside the region. Consequently, the results are clearly inconsistent with the limited-resource account. In contrast, the global-habituation theory would predict that a lower overall distractor frequency results in greater distractor interference for all locations. Accordingly, the pattern of results would be in line with a small global habituation effect (in addition to a local effect), which we may not have had enough power to resolve. However, this pattern could also be explained by an effect which, albeit being ‘local’, is somewhat fuzzy, ‘spreading’ to nearby (but not further-away) locations. This alternative explanation would predict that there would be a larger dependence of the interference on the distractor frequency outside the region for distractors on the border of a region, compared to locations ‘internal’ to the region (the latter being surrounded on both sides by locations which also belong to the region). In more detail, the argument is that if more frequent appearance of the distractor at one location results in stronger suppression of distractors at that location, but also somewhat stronger suppression of distractors at nearby locations (‘spreading’), then the amount of suppression on the border of a region would depend on some weighted average of the local frequencies in the two regions. Consequently, with the same local frequency in two to-be-compared regions, locations on the border would be more suppressed if the global frequency, and therefore also the local frequency in the other region, is higher, because of a stronger contribution to the suppression from the ‘spreading’ from the other region.

Recall that, in our experiments, the items bordering a region were those that were immediately above and below the leftmost and rightmost items (on the middle circle; see Figure 2); distractors never appeared at the leftmost and rightmost locations and the regions above and below these locations had different distractor frequencies. Any difference between border and internal locations would be expected to be particularly pronounced for the comparison of distractor interference between the rare region in the 80/**20** condition of Experiment 1 and the frequent region in the **80**/20 condition of Experiment 3 (which both had the same, 10% local distractor frequency), with the distractor frequency in the respectively other region (i.e., the frequent region with a local distractor frequency of 40% in Experiment 1 and the rare region with a local distractor frequency of 2.5% in Experiment 3) differing by a factor of 16 between the conditions (see Figure 1). Thus, to test the alternative, ‘spreading’ hypothesis elaborated above, we performed a mixed-effect ANOVA on distractor interference in the 10% distractor-frequency regions (**80**/20 condition in Experiment 3 vs. 80/**20** condition of Experiment 1) as between-participant factor and distractor location within the region (i.e., internal vs. on the border) as within-participant factor; see Figure 9 for a plot of the data. The interaction turned out significant (*F*(1, 42) = 16.9, *p* < .001, 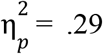, *BF*_incl_ = 143): as revealed by Bonferroni-corrected post-hoc *t*-tests, the interference was significantly smaller in the group with the higher global distractor frequency (the 80/**20** condition in Experiment 1) for locations on the border (56 ms difference, *t*(42) = 2.80, *p*_bonf_ < .05), but not for internal locations (−17 ms difference, *t*(42) = −0.86, *p*_bonf_ = 1), in a region. This supports the hypothesis that distractor-location learning effects are relatively local, though with some spreading to nearby locations.

**Figure 9:**
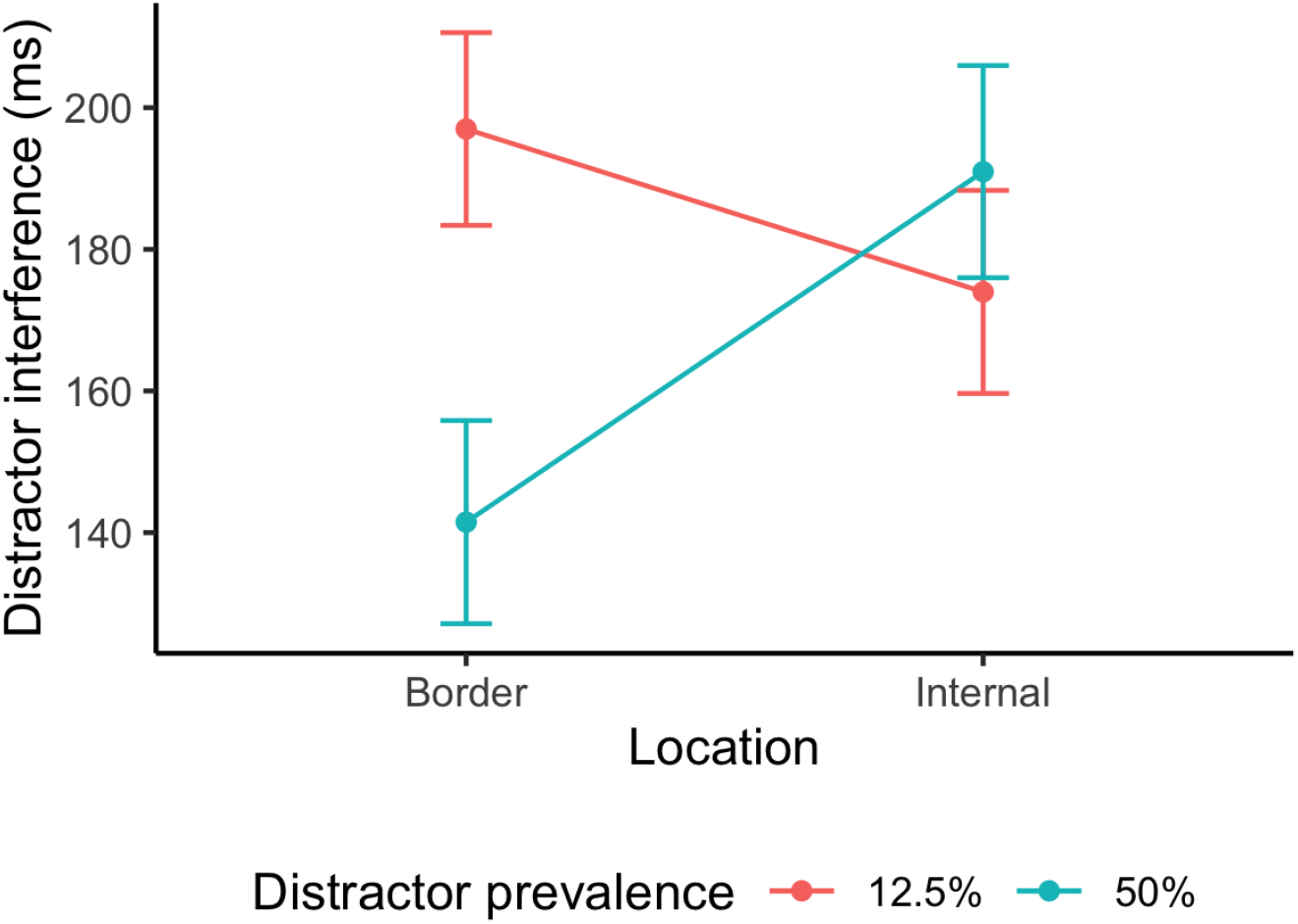
Distractor interference in two conditions in which a distractor appeared in a region with a 10% local distractor frequency, separated into trials on which a distractor appeared on the border of the respective region or internally within the region. The 50% distractor prevalence condition is from the rare region in the 80/**20** group of Experiment 1 and the condition with 12.5% distractor prevalence is from the frequent region in (**80**/20) Experiment 3. Error bars indicate the standard error of the mean.

## Model

Since our results are consistent with the distractor interference depending only on the local distractor frequency in a region, and not on how often the distractor appeared outside that region (at least when considering locations that do not fall on the border of a region), we went on to examine for the functional relationship between local distractor frequency and the amount of distractor interference to account for the results across all experimental conditions. To this end, we considered the amount of interference caused by distractors appearing in each region, in each of our experiments, as a function of the *local* distractor frequency in that region, and we fit several models to this combined dataset, representing different ways in which the amount of distractor interference could depend on local distractor frequency. Given that locations on the border of a region are influenced by carry-over effects from the other region (see “Within-region analysis” above), we based the model fitting on locations internal to each region. Figure 10 shows the combined data set from all our experiments and the predictions of the best-fitting model (see Supplementary for the full model comparison). The best model – which explains the amount of distractor interference quite well in all conditions (it somewhat underestimates interference for the lowest distractor frequency) – assumes that the interference grows linearly with the ‘Shannon information’ associated with a distractor. In more detail, this model assumes that the amount of interference depends on how much information is gained by observing a distractor at a particular location (or region), as formalized by Claude Shannon (1948). In essence, the Shannon information quantifies how much is learned from acquiring some new knowledge, quantified in terms of the number of two-alternative (e.g., yes/no) questions that had to be answered for gaining that knowledge. One might expect distractors which carry more information, in Shannon’s sense, to capture more attention, whereas a distractor that is expected based on stimulus history and thus relatively uninformative (or ‘unsurprising’) would be less likely to summon attention.^2^ While there are various ways in which distractor interference (*I*) could depend on the Shannon information (which are elaborated and tested in the Appendix), the winning model simply assumes that it is through a linear function:

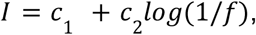

where *log(1/f)* is the Shannon information associated with a distractor occurring in a region with local distractor frequency *f*, and two parameters *c*_1_ and *c*_2_. The parameter *c*_1_ = 53 ms, indicates the lower asymptotic level of distractor interference if the distractor occured in the same region on every trial (i.e. 100% distractor prevalence and all distractors occur in the same region). The parameter *c*_2_ = 48 ms determines the rate of decrease of distractor interference with increasing local frequency. Of note, this model (with different parameter values) also provides a good fit to the distractor-interference effects when these are normalized to the respective distractor-absent RTs, in order to compensate for any general differences among the independent participant groups in response speed on the common baseline trials (see Appendix 3 for details).

**Figure 10:**
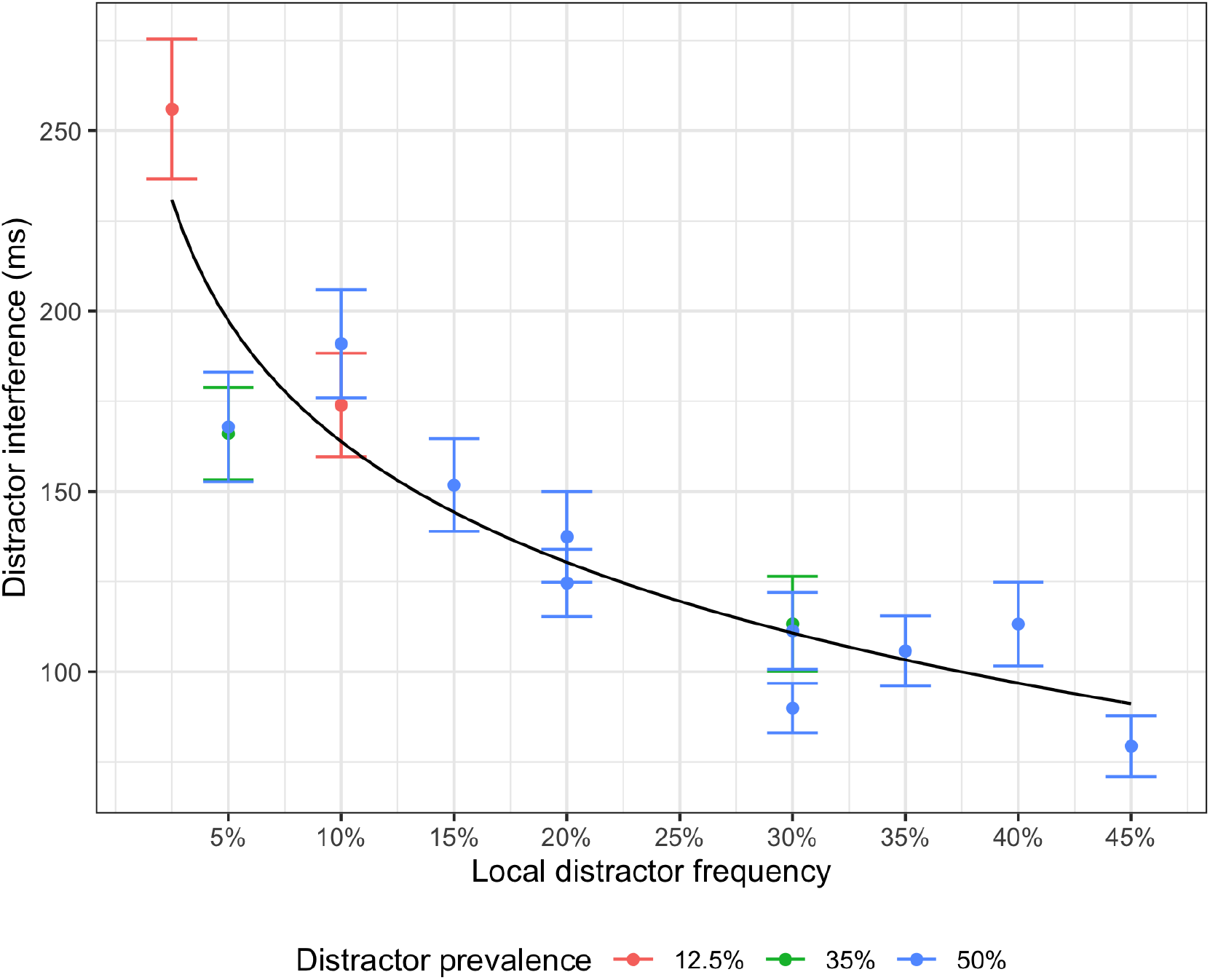
Distractor interference as a function of the local distractor frequency in a region, for distractors appearing at an internal location in in (i.e., not on the border of) the region, for all experimental conditions. Note that each experimental condition contributes two data points, one for trials with a distractors in the frequent and one for one for trials with a distractors in the rare region in that condition. The black line depicts the predictions of the best model: *I* = 53 + 48 · *log*(1/*f*). Error bars indicate the standard error of the mean.

## Discussion

In this study, we examined how the amount of response-time interference caused by a salient singleton distractor in a visual search task depends on the local distractor frequency in the region where the distractor appeared, as well as the frequency with which distractors appeared elsewhere in the search display. In Experiment 1, we found that across four conditions (tested in different groups of participants) with the same global distractor frequency, but differing ratios between the frequency of distractors in the (statistically) frequent and rare distractor region, the probability-cueing effect, that is, the difference in RTs between trials on which a distractor appeared in the rare compared to the frequent region, grew larger with increasing probability ratio (see Figure 3). A closely related result was recently reported by Lin, Li, Wang, and Theeuwes (2021), who used a similar paradigm, although with a single frequent distractor location (rather than a frequent region encompassing multiple locations) and a distractor defined in a different feature dimension to the target (see Footnote 1 for details). Despite some differences in the precise result pattern (see Footnotes 2 and 3 for details), taking Lin et al.’s effect of the probability-ratio (distractor at likely:unlikely location) on distractor interference together with effect of our distractor-region manipulation increases the confidence in the general result pattern revealed in Experiment 1.

Separate analyses (in Experiment 1) of distractor interference in the frequent and rare regions were somewhat inconclusive, but suggested that, if anything, the increased probability cueing-effect with increasing bias arises more from increased distractor interference with decreasing distractor frequency in the rare region, rather than decreasing distractor interference with increasing distractor frequency in the frequent region.^3^ Such an asymmetric pattern would be expected as response speed for the frequent region approaches the (distractor-absent RT) ceiling.

Interestingly, we also found a corresponding increase in the target-location effect (target in the rare minus target in the frequent distractor region) with increasing probability ratio, even though the target appeared equally frequently in both regions (see Figure 4), that is: RTs were slowed more when the target appeared in the frequent compared to the rare distractor region, the more the distractor frequency differed between these regions.^4^ This effect is consistent with the mechanism responsible for the distractor-frequency-dependent suppression operating on some (level of) spatial representation, or map, common to both the distractors and targets in our study. We further consider which representation this may be below.

A combined analysis of distractor interference indicated that the interference depended logarithmically on the local distractor frequency (see Figure 10), that is: interference decreased more rapidly with increasing distractor frequency when the frequency was low compared to when it was higher. This would explain further why we only found a significant difference in interference among the different spatial bias groups in Experiment 1 for distractors that occurred in the rare region.

In Experiment 2, when the local distractor frequency in the frequent distractor region was fixed, varying the distractor frequency in the rare region did not significantly influence the amount of interference caused by distractors in the frequent region, which is opposite to the prediction of the limited-resource theory (cf. Shaw & Shaw, 1977). In Experiment 3, when the local distractor frequency in one region was the same across two groups, the amount of distractor interference in that region did not significantly depend on whether this region had a four times higher or four times lower distractor frequency compared to the other region (see Figure 8). The limited-resource theory predicts that a lower distractor frequency in one region should result in less distractor interference in the other region, since more of the limited resource can be allocated to that region. Consequently, our results clearly contradict the limited-resource theory. On the other hand, the results are consistent with local habituation (Rankin et al., 2009; Sokolov, 1963; Turatto, Bonetti, & Pascucci, 2018), since local habituation predicts that the amount of distractor interference in a region should only depend on the local frequency of distractors in that region.

When further analyzing this effect separately for distractors on the border and distractors in the central part of the region, we found that for distractors falling on the border, but not in the central part, of a region, interference was significantly higher in the condition with a higher distractor frequency in the respectively other region. This suggests that while the effect is not global, it is also not so local as to be specific to an individual location in the search display. Rather, it can spread to locations at least two steps apart on the virtual display circle. (Recall that the frequent and rare distractor regions were, on each side, separated by a ‘neutral’ location where distractors never appeared and which therefore did not belong to either region; so, the items on the border of different regions were two steps apart.)

Combining the data from all experiments, we found the amount of distractor interference with RT to exhibit a non-linear dependence on the local distractor frequency, which could be explained by a model assuming that distractor interference has linear dependence on the amount of Shannon information associated with the local distractor probability. This is broadly consistent with the model of Itti and Baldi (2009), according to which ‘Bayesian surprise’ is a major factor in determining how likely a stimulus is to attract attention. Bayesian surprise quantifies how much an observer’s probabilistic model of the world is changed by a new observation. Focusing attention on stimuli with high Bayesian surprise makes sense from the theoretical view of ‘predictive coding’ (e.g., Clark, 2013; Friston & Kiebel, 2009; Rao & Ballard, 1999), since high surprise indicates that predictions are in need of updating, while for a stimulus resulting in low surprise predictions are already reliable. Unlike Shannon information, Bayesian surprise takes into account observers’ prior beliefs. This would be important for modeling how the distractor probability distribution is learned to begin with, as well as for explaining the results of studies in which the probabilities change substantially during an experiment (e.g., Valsecchi & Turatto, 2021). But it is arguably less important here, since we focused on performance after learning a distribution (recall that we excluded the first trial block in the analysis) that remains constant throughout the experiment.

Thus, while these results contradict the limited-resource theory, they are, in principle, consistent with a habituation account (Poon & Young, 2006; e.g., Rankin et al., 2009; Sokolov, 1963), with relatively location-specific habituation. That is, as participants are repeatedly exposed to a distractor in a particular region, the attentional orienting response triggered by distractors appearing in that region decreases, resulting in less attentional capture by salient stimuli in that region. With regard to our findings and their interpretation in terms of ‘local habituation’, two questions arise which we will discuss in turn: (i) Can our findings be reconciled with a recent report by Valsecchi and Turatto (2021), according to which distractor-location probability cueing involves a combination of both local and global factors? And (ii), is our finding that distractor-location learning is essentially based on the local distractor probability attributable to ‘habituation’ as “a non-associative or *task-free* learning mechanism that reduces neural responses to stimuli based on passive exposure”, or is it top-down mediated in the sense that “[the observer] must know what features *belong to distractors before* they can be suppressed” (Won & Geng, 2020, p. 1987)? Related to this, at what level in the functional architecture of search guidance is acquired local distractor-location suppression implemented?

### Role of global distractor frequency in distractor-location probability cueing?

Our proposal that it is essentially the local distractor probability that drives the distractor-location cueing effect appears to be challenged by an intriguing finding reported by Valsecchi and Turatto (2021), which points to the cueing effect also involving a global component. In Valsecchi and Turatto’s (2021) (online) Experiment 1, observers started with three ‘training’ blocks, with one high (40%), one intermediate (20%), and one low-frequency (6.6%) distractor location, which were equidistantly arranged around a ring consisting of a total of 12 locations (33.3% distractor-absent trials). Targets, too, could only occur at the three distractor locations, all other locations being occupied by non-target fillers. Distractors were salient color singletons, targets shape singletons. The results revealed a graded probability-cueing effect, that is, less interference by distractors at the high- vs. the intermediate- vs. the low-probability location, and a mirror image target-location effect on distractor-absent trials. The training blocks were followed by ‘extinction’ blocks without any distractors. In the subsequent two ‘test’ blocks, distractors were reintroduced, but now they appeared with equal probability at all three locations – importantly, with the same local probability as for the previous low-probability location (6.6%); that is, compared to the training blocks, the local probability was reduced for the previously high- and intermediate-probability locations, and, associated with, the global distractor probability. Valsecchi and Turatto found that the probability-cueing effect remained the same in the test blocks (compared to the last two training blocks), but distractor interference was overall increased, even at the location of previously low-probability distractor (at which the local distractor probability had not changed). They concluded that the overall increased distractor interference must result from the reduced global distractor probability (due to the reduction of the local probability at the previous high- and intermediate-probability locations)^5^.

Even though Valsecchi and Turatto’s experimental design was very different to ours, their results appear to be at odds with our failure to a significant influence of the global distractor probability on the probability-cueing effect. However, a few points about their study are noteworthy. First of all, during the learning phase, the variation of distractor probability (40%, 20%, 6.6%) would have resulted in a strong covariation of target probability: 20%, 35%, and 45% for the high-, intermediate-, and low-probability distractor location on distractor-present trials^6^, providing the potential for a confounding of statistical distractor-location with target-location learning. This may have overall inflated the ‘distractor-interference’ effects, which ranged from over 100 ms (high-probability location) to over 200 ms (low-probability location). These effects greatly exceed those typically reported in the additional-singleton paradigm (e.g., even in the online experiment of Lin et al., 2020, the effects were only about half for the 6:1 probability-ratio condition, which corresponds to the 40/6.6 ratio in Valsecchi & Turatto, 2021) – and they might have been responsible for the lack of ‘extinction’ in their study (evidenced by an undiminished local probability-cueing effect in the test blocks vs. the last two training blocks).

Nevertheless, it remains that the distraction effect was generally increased in the test blocks. This may be attributable to the generally increased ‘surprise’ value of any distractor appearing during the test blocks (especially after the extinction blocks without any distractors) – which gave rise to a delay in dealing with distraction, independently of where it arose (the previously high-, intermediate-, or low-probability location). Of note, the search RTs in Valsecchi and Turatto (2021) study were generally quite slow, with search very likely involving eye movements; and the size of their distraction effects (100–200 ms) would be indicative of oculomotor capture by the distractors on a substantial proportion of trials. It could therefore be that the generally increased distraction effect following the reduction of the global distractor probability is not due to a modulation of (oculomotor) capture itself, but to the time taken to disengage attention from a distractor that captured the eye – consistent with Sauter et al. (2020), who found oculomotor disengagement, measured in terms of the fixational dwell time on a distractor, to be faster from distractors at likely relative to unlikely display locations. For instance, when distractors become globally infrequent, it might take simply longer to decide that the item that captured attention is a distractor rather than the target, because the starting point in the decision (e.g., a stochastic diffusion) process shifts towards the ‘target’ and away from the ‘distractor’ boundary (Allenmark et al., 2018; for oculomotor evidence of such adjustments of post-selective decision criteria, see Allenmark et al., 2021). Thus, while massive reductions in the global distractor probability (such as from 66.6% to 33.3% in Valsecchi & Turatto, 2021) may indeed give rise to ‘global’ RT distractor-interference effects, these effects may be very different in nature (e.g., arising from post-selective item identification) to what we have described above as ‘global habituation’ (which would impact attentional capture directly). – In any case, in our study, there was never any change in the global (or local) distractor probability over the course of an experiment, over which participants also sampled the biased distractor distributions over 1440 trials (which compares with just 315 ‘training’ trials in Valsecchi & Turatto, 2021). Under these conditions, we found no compelling evidence of the global distractor probability (which varied substantially among our conditions, from 50% to 12.5%) driving the interference pattern.

Given that distractor-location learning is essentially ‘local’, two interrelated questions arise, namely: (i) is it to be considered a ‘local habituation’ process, and (ii) where, in the functional architecture of search guidance, does it occur? One defining property of ‘habituation’ is that it is “a non-associative or *task-free* learning mechanism that reduces neural responses to stimuli based on passive exposure”. This contrasts with “A core supposition of both [top-down accounts or statistical-learning accounts of distractor suppression] … that distractor suppression occurs only after the observer has knowledge, explicit or implicit, of what defines a distractor within the current context. These theories imply that one must know what features *belong to distractors before* they can be suppressed” (Won & Geng, 2020, p. 1987). Accordingly, while both types of theory might converge on where, in the cognitive architecture, the learning is implemented, they differ radically with regard to the cognitive dynamics assumed to underlie the learning. Based on an fMRI study (Zhang et al., 2021; as well as drawing on ideas from the oculomotor-capture study of Sauter et al., 2020), we have recently advocated an – in Won and Geng’s (2020) terms – active, ‘top-down’ view of statistical distractor-location probability learning, where the learning changes the responsivity of local feature-coding mechanisms in early visual cortex. We go on to describe this view, and how it explains the present data, in greater detail, before discussing possible challenges from passive ‘habituation’ accounts.

### Locus and dynamics of distractor-location learning (Zhang et al., 2021)

In Zhang et al.’s (2021) fMRI study, we used (in one condition) essentially the same stimuli as in the present experiments and a fixed distractor-probability ratio of 4:1 between the frequent and rare distractor regions. The results revealed distractor-generated signaling (indexed by the beta values) to be reduced in early visual-cortex areas – from V1 to V4 – for the frequent vs. the rare region, with the beta-values of distractors at (retinotopically mapped) locations in the frequent and, respectively, rare regions predicting the magnitude of distractor interference: the lower an individual’s beta value, the lower the interference.^7^ Consistent with standard notions of the functional architecture of visual search, we assumed that salient distractors may capture attention (prior to selection of the less salient target), by achieving a higher level of signaling on the search-guiding ‘attentional-priority’ map (e.g., Ferrante et al., 2018; Liesefeld & Müller, 2020; Müller et al., 2003; Wolfe, 2021). The distractors then need to be retroactively rejected (i.e., their priority signal needs to be suppressed) for attention to be disengaged from the distractor position and relocated to the target position – whose priority signal is initially weaker than that associated with the distractor, but stronger than that of the non-targets in the display (see also Sauter et al., 2020). Distractor suppression (on the priority map) then acts a feedback signal down to early visual areas, locally down-modulating the responsivity of the corresponding (entry-level) feature-coding mechanisms to the incoming perceptual signals. Thus, given that the stimulus-evoked activity is reduced as a result of ‘top-down tuning’ at early levels of signal coding, only a weakened signal is passed upwards in the processing hierarchy – ensuring that the signal strength (and thus the selection probability) of a spatially corresponding stimulus is reduced at the level of the search-guiding priority map. This statistical-learning dynamics would naturally explain not only why the representation of distractors becomes overall weaker if the global distractor frequency (or ‘prevalence’) is increased (e.g., Bogaerts et al., 2022; Geyer et al., 2008; Müller et al., 2009; Won et al., 2019), but also why it is increased more for regions in which distractors occur frequently vs. regions in which they occur only rarely. Further, it would explain why not only distractor signaling is reduced at the early visual coding level, but also target signaling – namely, because the top-down inhibitory feedback signals that down-modulate the responsivity of the early visual areas are relatively ‘feature-blind’ (“feature blindness” being considered a defining characteristic of purely spatial saliency and priority signals; see, e.g., Fecteau & Munoz, 2006). Critically, however, rejecting a distractor, and thus inhibiting its location from the priority map downwards, requires a (more or less explicit) process of identifying an attended distractor as an erroneously selected, task-irrelevant ‘non-target’ item. In the words of Zhang et al. (2021), “‘distractor’-location inhibition is top-down mediated” in that it is “tied to the status of distractors as ‘distractors’” (p. 13) – and this would be at variance with a strict notion of ‘local habituation’.

### Passive versus active learning of local distractor probabilities

However, there appear to be a number of reports in the literature that are at odds with this view of active, top-down-mediated distractor-location learning – Turatto, Bonetti, Pascucci, and Chelazzi (2018), Won and Geng (2020), and Duncan and Theeuwes (2020) –, which we discuss in turn.

*Turatto et al. (2018)* asked their observers (in their Experiment 2) to passively view a series of displays (each consisting of four placeholder circles), 50% of which included a salient, luminance-onset stimulus (superimposed on one of the circles). When the salient stimulus later became a distractor during a target discrimination task (in which a precue, presented prior to the briefly flashed distractor, indicated the circle in which the to-be-discriminated target would later appear), attentional capture was attenuated by some 50% (already in the initial block of the active task) compared to when the displays were not first passively viewed (in Experiment 1). The authors concluded that the passive exposure habituated the visual system to the salient stimulus, effectively reducing its attention-capturing power during a later active task. However, since Turatto et al. used identical stimulus displays during passive exposure and the active task, it remains possible that ‘surreptitious’ (but re-/active) suppression of the salient stimulus during passive viewing led to the reduction of distractor interference already in the active task. This may be so especially because the salient distractor was a high-luminance white annulus frame (52.5 cd/m^2^) briefly (for 100 ms) flashed against a dark-gray background (0.07 cd/m^2^) and highlighting one of four previewed placeholder circles (7 cd/m^2^). Such highly salient sudden-onset stimuli would have acted as strong attractors of eye movements, which were, however, to be prevented by instruction: “In the passive blocks …, the cue was presented but the target was not, and *participants were asked to maintain fixation on the central spot*, while passively viewing the display” (Turatto et al., 2018, p. 1832; our italics). In other words, the distractors would have attracted attention and the eye at least initially, and participants would have had to disengage attention from their locations, involving re-/active suppression, to reorient to the central fixation marker. That is, the task demands are similar to when a salient distractor is to be avoided in an active search task.

*Won and Geng (2020)* attempted to deal with this issue by introducing a novel two-phase paradigm. During the first, ‘training’ phase, observers in the critical ‘habituation’ group were passively exposed to one set of (heterogeneous) colors on four task-irrelevant ‘circles’ in the display periphery (no-task trials), while they actively searched for a (gray) target square (and reported a numeral inside it) presented among three heterogeneously colored ‘distractor’ squares in the display center (search-task trials). Importantly, no-task (circle-display) and search-task (square-display) trials were interleaved, and the sets of circle (‘new set’) and square colors (‘trained set’) were non-overlapping. During the second, ‘testing’ phase, a subset of the ‘habituated’ colors from the outer circles were introduced as ‘new colors’ of the central search distractors, randomly interleaved with the ‘trained set’ of colors. Search performance in this ‘habituation’ group was compared with a ‘control’ group (in Experiment 1) that, both during training and test, was exposed to non-colored circle displays (the circles were simple gray outline circles that were not color-filled-in); the search displays were identical to those in the ‘habituation’ group (i.e., in the test phase, the colors of the search distractors were ‘trained’ or ‘new’). The critical comparison concerned (what Won and Geng referred to as) ‘distractor interference’ on search trials in the test phase, calculated as the search RT to displays with the ‘new’ color distractors minus the RT to displays with ‘trained’ color distractors. The results revealed search to be slower for the new vs. the trained color distractors, that is, in Won ad Geng’s terms, there was ‘distractor interference’; and, importantly, the interference was significantly reduced for the habituation group compared to the control group. In other words, viewing irrelevant color items under no-task conditions (in the training phase) later on helped observers find the (gray) target faster when it was presented among distractors with these previously irrelevant colors, compared to observers who had no pre-exposure to these colors.

This indeed shows that pre-exposure to colors even under no-task conditions facilitates search when the same colors later on appear in the search distractors. Won and Geng’s (2020) preferred interpretation of this is that the new distractor colors introduced in the test phase had a higher ‘novelty’ (or surprise) value for observers in the habituation compared to the control group, as a result of which the (new-color) distractors may have engaged more attention upon search display onset, thus delaying target selection. On the other hand, though, what Won and Geng (2020) investigated was not interference caused by a salient singleton distractor, but rather the effect of multiple, color-heterogenous *non-targets* in a search scenario in which the target (even though it was defined by a unique color) may not have popped out. In classical pop-out search tasks, the non-target context facilitates target detection by increasing the target feature contrast while reducing the distractor feature contrast (e.g., Bravo & Nakayama, 1992; Liesefeld et al., 2016). In contrast, with the color-heterogenous distractors in Won and Geng’s scenario, all items would have had a relatively high color contrast (each item differed from the other in color), so that search would have been mainly template-based (Liesefeld & Müller, 2020), and the target template would have been less optimally tuned to the (variable or broader) non-target color context in the test phase when only a limited set of colors was encountered in the preceding training phase. Though, what Won and Geng’s findings show under this interpretation is that even search-irrelevant colors encountered in the training phase are integrated in the target template. But, what Won and Geng interpret as (bottom-up) non-target interference might also reflect the relative lack of (top-down) target facilitation.

Another point to note is that the statistical learning, or ‘habituation’, effects demonstrated by Won and Geng (2020), as well as to some extent those of Turatto et al. (2018)^8^, are non-spatial in nature: they reflect the learning of, or adaptation to, the features of more or less salient distractors (or non-targets) in visual search, but not the learning of their positions within the search displays. In Won and Geng (2020), the task-irrelevant colored circles were (deliberately) presented at variable locations further in the periphery compared to the central search displays (to make them appear unrelated to the search task); and in Turatto et al. (2018), the luminance-onset distractors occurred equally likely at all (four) possible display locations.

There is one recent study, however, that did examine passive learning of the spatial distribution of salient singleton-color distractors: *Duncan and Theeuwes (2020)*. The experiment consisted of two phases, with different tasks. In the first, ‘learning’ phase, participants had to judge, as fast and accurately as possible, whether the item arrangement (made up by eight colored shapes) formed a global diamond (made up of the local elements) or a global circle. All items were of one color, except – on 67% of the trials – for one singleton color item, which appeared more likely at one location (65% of trials), compared to any of the seven other locations (35%/7 = 5% of trials). Likely locations were restricted to the 12, 3, 6, and 9 o’clock locations, which were identical in the two types of global (diamond, circle) arrangement. Participants performed three blocks of this task. In the second, ‘test’ phase (blocks 4 and 5), the task was switched to the ‘usual’ search for a singleton shape target in the possible presence of an additional color singleton; the response required discrimination of the (horizontal, vertical) orientation of a line inside the shape target. Importantly, the additional color singleton now appeared equally likely at all eight display locations. The critical finding was that, despite the removal of the spatial distractor-location bias in the test phase, in block 4 (but no longer in block 5) RTs were faster when the distractor appeared at the previously high-, vs. a low-, probability location (significant effect in one-tailed testing in online Experiment 4, with 80, out of initially 120, participants who had not dropped out after the learning phase; non-significant in onsite Experiment 1 with 24 participants). This is consistent with participants having learned distractor-location probability distribution in phase 1 (even though the task required a global item-configuration judgment, rather than selection of any local item), which was then carried over to the search task where distractors occurring at the (previously) likely location caused initially less interference than distractors at one of the unlikely locations (and this cueing effect was then unlearned over the course of the search task due to the unbiased distractor distribution).

It is important to note, though, that when the carry-over effect was significant (in their Experiment 4), there was also slight, but highly significant distractor-location effect in the initial learning phase: the global-configuration judgments, although generally quite fast (compared to the compound-task response required in the search task), could be issued faster when the singleton color item appeared at the likely location, compared to any of the unlikely locations (and on both types of color-singleton-present trials, RTs were slower compared to -absent trials). In other words, it appears likely that the salient color singleton did interfere with performance of the global task (by summoning attention to the local level; e.g., Navon, 1977) and so may have been actively suppressed to mitigate the intrusion of a ‘disturbing’ distractor. In fact, this possibility is expressly acknowledged by Duncan and Theeuwes (2020), who state that “… while our results clearly show that SL [statistical learning] may occur in the absence of [goal-directed] top-down attention, we cannot rule out that bottom-up capture may still play a role” (p. 63).

Thus, we contend that prior studies (Duncan & Theeuwes, 2020; Turatto et al., 2018; Won & Geng, 2020) provide no compelling evidence against our view that distractor-*location* probability learning is an ‘active’ (though not necessarily explicit) process, that is: a process involving having to identify a distractor that captured attention as an erroneously selected item and to reject it (i.e., suppress its priority signal) in order to disengage attention and reorient it to the target item. In the learning phase of Duncan and Theeuwes’ (2020) study, this would mean to disengage attention from the local item level and reorient it to the global item configuration; in our task, to reorient it to the item with the second most high attentional priority. This retroactive, local distractor rejection/suppression provides the top-down feedback signal that down-modulates the responsivity of feature-coding neurons at the corresponding locations in early visual cortex, thus naturally implementing local statistical distractor-location learning. We contend that this process of distractor rejection is an ‘active’, top-down process, which, however, does not mean that it is a ‘willed’, intentionally explicit process (see also Gaspelin & Luck, 2018b). Rather, it is likely to operate implicitly as an, over the course of an experiment, automatized routine. This is evidenced by the fact that, although our participants collectively displayed some above-chance explicit knowledge of the distractor-location probability distribution, the cueing effects did not differ between observers who, in the explicit-recognition tests performed after the experiments, had correctly indicated the bias in the distribution and those who hadn’t.

## Conclusion and Outlook

In summary, distractor-location probability cueing is based largely (if not exclusively) on the local distractor frequency, rather than involving the (re-) distribution of a global inhibitory resource (in contrast to target-location probability cueing; cf. Shaw & Shaw, 1977). While we found no evidence of the global distractor prevalence playing a significant role in local learning, more work is required to understand the nature of modulatory ‘global’ effects reported in the literature (cf. Valsecchi & Turatto, 2021; see also Bogaerts et al., 2022; Müller et al., 2009; Won et al., 2019). Further, while we have firmly established distractor-location learning as an essentially ‘local’ phenomenon, it remains to be seen whether it is to be classed as ‘habituation’ in the strict sense of “attenuated processing of previously encountered sensory information that did not elicit attentive processing or a behavioral response” (Won & Geng, 2020, p. 1994) – as has been argued by some; or, alternatively, whether it is top-down mediated in that it results from observers identifying a distractor that captured attention as a ‘distractor’, as a precursor to (actively) suppressing its location and so releasing attention to be reoriented elsewhere in the service of search – as we have argued (e.g, Zhang et al., 2021).

## Appendix 1: Model comparison

Our results show that the amount of RT interference caused by a distractor does not significantly depend on the global frequency distribution of distractor locations, but only on the local frequency with which distractors occur within a given region. Accordingly, we combined the data from all our experimental conditions in order to examine the functional dependence of distractor interference on the local distractor frequency. Here, we compare a number of different models of how the interference could depend on the frequency. As a baseline against which to assess the various models, we included a simple linear model which assumes that distractor interference decreases linearly with increasing local distractor frequency:

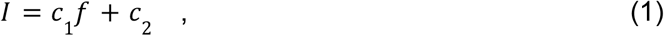

where *I* is the amount of distractor interference, *f* is the local distractor frequency, and *c_1_* and *c_2_* are free parameters of the model determining the slope and intercept of the linear fit.

Table S1 shows the difference between the Akaike Information Criterion (AIC; Akaike, 1974; Vrieze, 2012) of each model and the AIC of the linear model (more negative values indicate that the model performed better). We also include an exponentially decreasing function, based on the assumption that the amount of reduction of distractor interference resulting from an increase in distractor frequency might be proportional to the amount of interference before the increase:

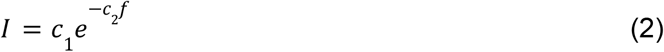

Next, it could be that the attentional priority, or ‘selection salience’ (Zehetleitner et al., 2013), of the distractor decays exponentially with the local distractor frequency, the probability of attentional capture is determined by the distractor salience normalized by the total salience (because the probabilities of selecting the different items in the search display should sum to one), and the RT distractor interference is proportional to the probability of attentional capture:

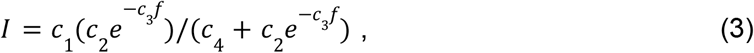

where *c_1_* represents the RT interference produced by the distractor and *c_4_* represents the total salience of the other items in the search display (other than the distractor).

For the remaining models, we considered the possibility that the amount of interference depends on how much information is gained by observing a distractor in a particular location/region, as formalized by Claude Shannon (1948). One might expect distractors which carry more information, in Shannon’s sense, to capture more attention, whereas a distractor that is expected based on stimulus history and therefore uninformative (or ‘unsurprising’) would be less likely to capture attention. We assume that the Shannon information associated with a distractor depends on the local frequency as log(1/f)^9^. The simplest way in which distractor interference could depend on the Shannon information is through a simple linear function:

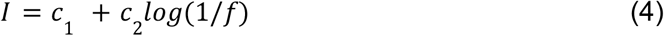

Alternatively, the distractor interference may increase exponentially with increasing Shannon information, this would predict a power-law dependence on the frequency:

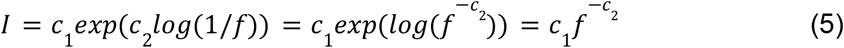

Finally, we considered the possibility that the salience of the distractor is proportional to the Shannon information and the distractor interference is determined by the normalized salience (similar to equation 3 above):

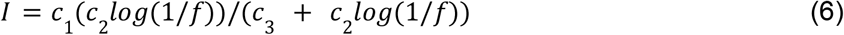

**Table S1:** The difference between the Akaike Information Criterion (AIC) of each model and the AIC of the linear model. More negative values indicate better model performance. The best model in terms of the AIC was the model which assumes that the distractor interference depends linearly on the Shannon information. The optimal parameters for this model were *c*_1_ = 53, indicating the predicted amount of distractor interference (53 ms) if the distractor occured in the same region on all trials, and *c*_2_ = 48, determining the rate of the non-linear decrease of distractor interference with increasing local frequency.

| Model | $\Delta\text{AIC}$ |
| --- | --- |
| Linear (eq. 1) | 0 |
| Exponential (eq. 2) | -2.7 |
| Normalized exponential (eq. 3) | 1.5 |
| Linear information (eq. 4) | -6.5 |
| Exponential information (eq. 5) | -6.0 |
| Normalized information (eq. 6) | -3.0 |

## Appendix 2: Re-analysis Lin, Li, Wang, & Theeuwes (2021)

Experiment 1 was similar in design to Lin et al. (2020), who however used a variation of Theeuwes’ (1992) standard ‘additional-singleton paradigm’. Their task required search for a shape-defined target singleton, in the potential presence of an ‘additional’ color-defined distractor singleton. Critically, a single location (out of 8 possible locations) was most likely to contain a distractor, whereas all other locations had the same, low distractor probability. The overall distractor probability averaged some 67% across the various ‘bias’ conditions (i.e., there were, on average, 33% distractor-absent trials), and the location-probability biases (ratio of single likely location to any of the unlikely locations) varied from 2:1 to 8:1. The effect pattern reported by Lin et al. (2020) was that distractor interference was generally reduced for the frequent vs. the rare distractor locations (i.e., there was a distractor-location probability-cueing effect), the more so the more likely the distractor appeared at the likely location (and the less likely it appeared at any of the unlikely locations). In principle, this pattern is consistent with two of the accounts considered in the Introduction: distribution of a limited inhibition resource and, respectively, local habituation.^10^

However, although Lin et al. (2021) implemented a similar probability-ratio manipulation to our Experiment 1, they did not analyze their data in quite the same way as we did. In particular, they did not report RT distractor interference separately for the single frequent distractor location and the rare locations, but only the difference between the two conditions (i.e., the distractor-location probability-cueing effect). Given this, here present our own analysis of their (freely available) data.

**Figure S1:**
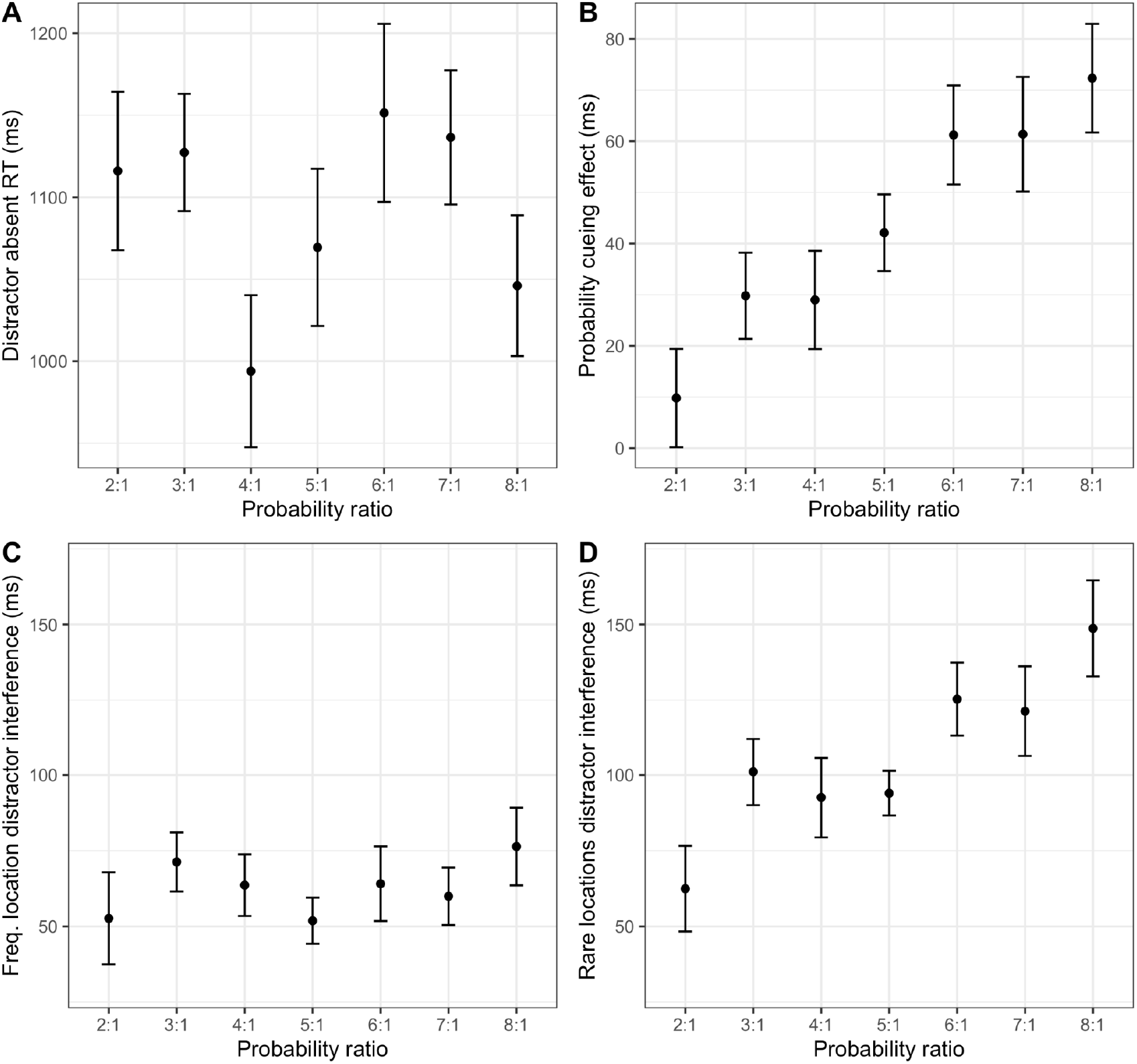
Distractor absent RTs (A); Probability-cueing effect (B); Interference by distractors at the single frequent location (C) and at the (average) rare location (D), in the data of Lin, Li, Wang, & Theeuwes (2021). Error bars indicate the standard error of the mean.

Figure S1 plots the distractor-absent RTs, the distractor-location probability-cueing effect, and distractor interference by distractors at the single frequent and, respectively, the (average) rare locations in the data of Lin et al. (2021). Similar to our Experiment 1, the increasing probability-cueing effect with increasing probability ratio results mainly from increasing interference by distractors occurring at one of the rare locations (where increasing probability ratio was accompanied by decreasing local distractor frequency). A mixed ANOVA revealed a significant effect of probability ratio on distractor interference caused by rare-location distractors (F(6,105) = 4.98, p < .001, 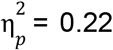). However, frequent-location distractors did not produce a decrease in interference with increasing probability ratio; if anything, there was a slight increasing trend (see Figure S1 panel C), though this was not statistically significant (F(6,105) = 0.68, p = .66).

## Appendix 3: Model fit and re-analysis with normalized data

Since we observed numerical differences in the distractor-absent RTs among the various conditions (i.e., independent-participants groups) which, while statistically non-significant, were not completely negligible in magnitude, we examined whether we would obtain similar results if we compensated for these differences in general response speed by normalizing the distractor interference with reference to the distractor-absent RTs. That is, for each participant, we divided the distractor interference by their distractor-absent RT, and we repeated the analysis described in Appendix 1 (and the Model section) with the resulting normalized distractor-interference score. As can be seen from Figure S2, we obtained very similar results, and the same model (though with different parameter values) still provided a good fit to the data.

**Figure S2:**
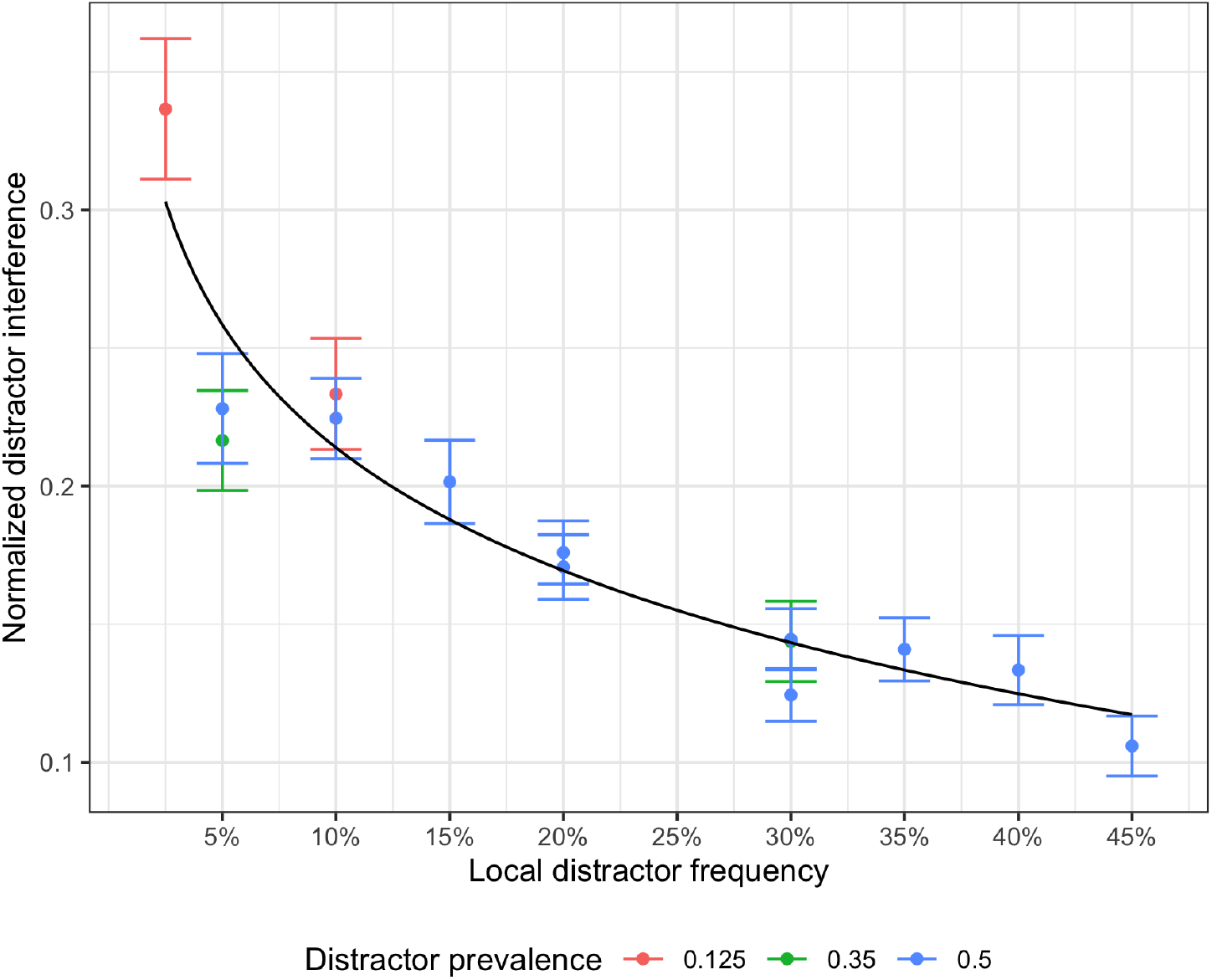
Normalized distractor interference as a function of the local distractor frequency in a region, for distractors appearing at an internal location in (i.e., not on the border of) the region, for all experimental conditions. Note that each experimental condition contributes two data points, one for trials with a distractor in the frequent and one for one for trials with a distractor in the rare region in that condition. The black line depicts the predictions of the best model. Error bars indicate the standard error of the mean.

In addition, below, we also repeat critical comparisons between conditions with the same local frequency in a region, but a different global distribution (for which the local and global theories make different predictions), based on normalized distractor interference from trials with a distractor at a region-internal location.

Based on Experiment 2, there are three such comparisons (see Figure 1), two comparisons to the 60/40 condition (onsite experiment and online replication) and one comparison to the 90/10 condition. The normalized distractor interference (for internal distractor locations) did not differ significantly between distractors in the frequent region in the reduced distractor-frequency condition of Exp. 2 and the frequent region in the 60/40 condition of Exp. 1, with the Bayes factor favoring the null hypothesis (–1-ms difference, t(39.2) = -0.042, p = .97, BF_10_ = 0.30). There was also no significant difference compared to the online replication of the 60/40 condition (19-ms difference, t(36.3) = 1.1, p = .25, BF_10_ = 0.50). Finally, the normalized distractor interference did not differ significantly between distractors in the rare region in the reduced distractor frequency condition of Exp. 2 and the rare region in the 90/10 condition of Exp. 1, with the Bayes factor favoring the null hypothesis (–12-ms difference, t(41.7) = - 0.44, p = .66, BF_10_ = 0.32).

We also considered the comparison between normalized distractor interference by distractors in the frequent region of Exp. 3 and in the rare region in the 80/20 condition of Exp. 1, which also did not differ significantly, with the Bayes factor favoring the null hypothesis (9-ms difference, t(38.3) = 0.36, p = .72, BF_10_ = 0.32).

Overall, the re-analysis of the critical (equal local distractor frequency but different global distribution) comparisons, based on normalized distractor interference and distractors appearing at region-internal locations, clearly favors the local learning hypothesis.

## Footnotes

1 In this regard, Experiment 1 is similar in design to Lin et al. (2020), who however used a variation of Theeuwes’ (1992) standard ‘additional-singleton paradigm (i.e., search for a shape-defined target singleton, in the potential presence of an ‘additional’ color-defined distractor singleton), in which – critically – a single location (out of 8 possible locations) was most likely to contain a distractor, whereas all other locations had the same, low distractor probability. However, as we have pointed out elsewhere, there is a potential problem with the ‘single-likely-distractor-location’ paradigm, namely: co-variation of *distractor*-location probability with *target*-location probability, making it difficult to determine the interference effects caused by distractors at the likely or, respectively, an unlikely location unconfounded by target-location learning. For this reason, we prefer the region-cueing paradigm, that avoids such confounding. For more details, as well as a reanalysis of Lin et al.’s data, see Appendix 2.

2 Mathematically, counting the yes/no questions corresponds to taking the base-two logarithm of the number of (equally probable) alternatives. This is the same as taking the logarithm of 1/p, where p is the probability of each alternative. This formula can also be used when there are no equally probable alternatives. In our case, if we take the probability of the distractor appearing in a region to equal the frequency with which it appears in the region in our experiment, then the log(1/f) in our model is the Shannon information associated with observing a distractor (in the region in which it occurs with frequency f).

3 Of note, in Lin et al.’s (2021) experiment, the increase in the distractor-location (probability-cueing) effect with increasing probability ratio between distractors at the likely vs. an unlikely location appeared to be associated with a general increase in distractor interference with the probability ratio, not only for unlikely locations (as in the present Experiment 1), but, if anything, also for likely locations (which, in the present Experiment 1, showed a decrease). See Appendix 2 for our re-analysis of distractor interference in the rare and frequent distractor locations in the experiment of Lin et al. (2021), who provided only an analysis of the difference between the two conditions (the cueing effect), but not of the underlying conditions themselves. This lack of a trade-off of distractor interference between the likely and unlikely locations makes it difficult to interpret the change in the cueing effect with increasing ratio theoretically, in terms of either a limited-inhibition-resource or a local-habituation account.

4 The equivalent effect was not significant in the study of Lin et al. (2021): while there was a main effect of target-location (i.e., the target-location effect differed significantly from zero), it was not significantly modulated by the (distractor at likely:unlikely location) probability ratio (*F*(6, 105) = 0.43, *p* = .856, 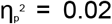, *BF*_10_ = 0.06). Such a modulation would have been expected if the mechanism responsible for distractor-frequency-dependent spatial suppression operates in a manner independent of the particular (in Lin et al.’s study: color and shape) features that single out the distractor and target, respectively.

5 A similar result pattern was obtained in their Experiment 2, which omitted the extinction blocks between the training and test blocks.

6 This calculation assumes that with a distractor at one location, the target was equally likely to appear at the two other locations.

7 With search for an orientation-defined target, this relationship was not obtained for distractors defined in the color dimension, which we attributed to a later, color-based distractor-filtering stage (involving the fusiform gyrus; see Zhang et al., 2021, for details).

8 In their study, the distribution of distractors was spatially uniform and distractors only occurred at a fixed set of locations at which, in the active task, targets would appear, too (on different trials, as the distractor and target locations never coincided).

9 That is, for simplicity, we assume that the probability of observing a distractor (in a particular location) in the definition of Shannon information is simply the local distractor frequency. This cannot be exactly right, since it predicts that the information associated with a distractor goes to infinity as the frequency approaches zero, but it may be a good approximation across our range of distractor frequencies.

10 Of note, though, there is a potential problem with the ‘single-likely-distractor-location’ paradigm: given that the distractor never occurs at the target location, the likely distractor location would necessarily be less likely to be the target location – the more so, the more likely the distractor occurs at the ‘likely’ location (i.e., more so in the 8:1 than in the 2:1 ratio condition). This imbalance would be exacerbated if the probability of distractor-absent trials (on which the target appears equally likely at all locations) is low (only some 33% in Lin et al’s study, compared to more typical 50%). That is, it becomes difficult to rule out the confounding of a *distractor*-location probability-cueing effect by a *target*-location probability-cueing effect. There is a way to correct for the reduced target probability at the likely distractor location, namely (as proposed, e.g., by Zhang et al., 2019): by compensatorily increasing the probability of the target appearing at the likely distractor location on trials on which the distractor appears at an unlikely location. In fact, a reanalysis of Lin et al.’s (2020) data shows that they did apply this correction. However, this again is problematic in that, especially with increasing ratios (e.g., from 2:1 to 8:1) of distractors occurring at the likely vs. an unlikely location, the actual occurrence of a distractor at an unlikely location would become increasingly predictive of the target appearing at the likely location. These contingencies make it difficult to determine the interference effects caused by distractors at the likely or, respectively, an unlikely location unconfounded by target-location learning.

## Notes

Author note: This study was supported by German Science Foundation (DFG) research grants MU773/16-2, awarded to HJM and ZS, and MU773/14-2, awarded to HJM. This study was not preregistered. The data and modeling code are available in the following GitHub repository: https://github.com/msenselab/Local_Inhibition

### Competing Interest Statement

The authors have declared no competing interest.

